# Optogenetic actin network assembly on lipid bilayer uncovers the network density-dependent functions of actin-binding proteins

**DOI:** 10.1101/2024.12.25.630233

**Authors:** Kei Yamamoto, Makito Miyazaki

## Abstract

The actin cytoskeleton forms a mesh-like network that drives cellular deformations. The network property is defined by the network density and the species of actin-binding proteins. However, the relationship between the network density, the penetration ability of actin-binding proteins into the network, and resulting network dynamics remains elusive. Here, we report an *in vitro* optogenetic system, named OptoVCA, which induces Arp2/3 complex-mediated actin network assembly on a lipid membrane. By changing the illumination power, duration, and pattern, the OptoVCA flexibly manipulates the density, thickness, and shape of the actin network. Taking these advantages, we examine the effects of the network density on two representative actin-binding proteins, myosin and ADF/cofilin. We find that the penetration of myosin filaments into the network is strictly inhibited by only a several-fold increase in network density due to the steric hindrance. Furthermore, penetrated myosin filaments induce directional actin flow when the network has a density gradient. On the other hand, ADF/cofilin penetrates into the network regardless of network density. However, network disassembly is dramatically inhibited by only a several-fold increase in network density. Thus, the OptoVCA contributes to understanding cell mechanics by examining the network density-dependent effects on actin-binding proteins.

## Introduction

The mesh-like actin network is a ubiquitous structure of the actin cytoskeleton, playing pivotal roles in dynamic cellular deformations such as a lamellipodium, bleb, and contractile ring (Kadzik, Homa, and Kovar 2020; García-Arcos et al. 2024). The network properties that characterize the specific subcellular structures are determined through interactions between actin filaments (F-actin) and actin-binding proteins (ABPs) related to each structure (Svitkina 2018; Blanchoin et al. 2014; Clarke and Martin 2021; Pollard 2016). To initiate the actin network assembly, actin polymerization is triggered by various nucleation-promoting factors (NPFs). The Wiskott-Aldrich syndrome protein (WASP) and WASP-family verprolin-homologous protein (WAVE) are the two major NPFs that activate a nucleator, the Arp2/3 complex, which in turn initiates branching of F-actin (Kurisu and Takenawa 2009; Derivery and Gautreau 2010; Firat-Karalar and Welch 2011). These NPFs are activated on lipid membranes including plasma membranes and organelle surfaces, and control the actin network assembly (Alekhina, Burstein, and Billadeau 2017; Breitsprecher and Goode 2013). During and after the actin network assembly, various ABPs interact with the network. Among ABPs, myosin II is a primary force generator that induces cellular deformations by generating contractile force with F-actin (Vicente-Manzanares et al. 2009; Craig and Woodhead 2006). ADF/cofilin (hereafter referred to as cofilin) disassembles the actin network by severing actin filaments, releasing the mechanical stress and enabling the dynamic recycling of F-actin (Bravo-Cordero et al. 2013).

Recent studies suggest that not only the species of ABPs but also tight regulation of the actin network density is important to achieve cellular functions. For example, it has been reported that myosin motors cannot generate proper contractile force both in sparse and dense actin networks in mammalian culture cells (Chugh et al. 2017). In addition, it has been reported that the penetration ability of ABPs into the actin network is affected by their protein sizes; smaller proteins such as ERM family proteins can enter the network, while larger proteins such as myosin filaments are frequently observed at the interior side of the cortical actin network (Truong Quang et al. 2021). Another potential function of the actin network is to serve as a steric barrier for proper organelle distributions in the cytoplasm (Moore et al. 2021; Olguin-Olguin et al. 2021). This growing evidence supports the idea that the actin network density is one of the key factors that broadly regulates not only the cell mechanics but also cellular functions mediated by ABPs and organelles. However, it is technically difficult to address the relationship between the protein sizes, the actin network densities, and the resulting functions of ABPs using live cells because of the complexity, thinness, and dynamicity of the actin networks (Laplaud et al. 2021; Jawahar et al. 2024).

To understand how the interactions between the actin network and ABPs are self-organized into specific structures, recent efforts have been devoted to reconstituting the actin cytoskeleton *in vitro*. The patterned illumination of UV light to a PEG-coated glass enables the binding of NPFs in the specified regions (Boujemaa-Paterski et al. 2017; Manhart et al. 2019; Bieling et al. 2016; Anne-Cécile Reymann et al. 2010, 2012). These studies have elucidated the characteristics of actin networks resulting from the various distributions and densities of NPFs on the substrates. Recently, the techniques to bind NPFs on a spatially patterned lipid bilayer have been developed, making it possible to analyze actin dynamics in more biologically relevant conditions (Colin et al. 2023; Yamazaki et al. 2024). However, it is still technically challenging to control actin polymerization in both space and time. Moreover, it is also difficult to assemble actin networks with various densities on the same lipid surface. Hence, further flexibility for the control of actin polymerization in the reconstituted systems has been desired.

To achieve this, optogenetic approaches offer great potential for manipulating actin cytoskeletal dynamics. Light-induced dimerization is a powerful method for spatiotemporal control of the intracellular localization of target proteins, and among these systems, iLID-SspB has been widely used (Guntas et al. 2015; Yamamoto et al. 2021; Mahlandt et al. 2023; Meiring et al. 2022; Martínez-Ara et al. 2022; Nakamura et al. 2023). Furthermore, iLID-SspB can be easily purified, making it well-suited for integration with *in vitro* experiments (Di Iorio et al. 2023; Matsubayashi et al. 2024). Here, we develop an iLID-SspB-based *in vitro* optogenetic system that allows us to control the actin network densities, thicknesses, and shapes with high temporal and spatial resolutions on a lipid bilayer. First, as a proof of concept, we use mammalian culture cells and show that the actin cortex is formed by light-induced recruitment of the VCA domain of WAVE1 to the plasma membrane. Actin polymerization induced by our optogenetic system, named OptoVCA, is dependent on the Arp2/3 complex. Next, we expand this strategy for *in vitro* experiments by combining OptoVCA with a mixture of purified actin cytoskeletal proteins and a supported lipid bilayer (SLB). Using this system, we demonstrate that the penetration ability of ABPs into the actin network and functions of ABPs are tightly regulated by the network density and the size of ABPs. These observations highlight the importance of the actin network density to understand cell mechanics.

## Results

### Optogenetic manipulation of actin polymerization in live cells

To control the assembly of actin networks by light illumination, we used WAVE1, one of the major NPFs in mammalian cells. WAVE1 contains the V domain and CA domain, which interact with G-actin and the Arp2/3 complex, respectively. These domains are collectively called the VCA domain (Kurisu and Takenawa 2009) (Fig. 1a). WAVE1 is inactivated by the WAVE regulatory complex (WRC). When WRC is recruited to the plasma membrane through the interactions with Rac1, PIP_3_, and IRSp53, the VCA domain is exposed to the cytoplasm. The activated WAVE1 recruits G-actin and the Arp2/3 complex to the VCA domain and promotes actin nucleation, leading to actin polymerization (Kurisu and Takenawa 2009). In line with this molecular mechanism, it has been reported that artificially localized VCA domain to the actin cortex induces cortical thickening in mouse oocytes (Chaigne et al. 2015). Light-induced oligomerization or membrane targeting of NPFs also enables control of actin polymerization (Nakamura et al. 2023; Taslimi et al. 2014). Therefore, we expected that a light-induced local increase in the density of the VCA domain would be sufficient for promoting actin polymerization both *in vivo* and *in vitro*. Particularly, two VCA molecules are required to activate the Arp2/3 complex (Yamaguchi et al. 2002), suggesting that the basal Arp2/3- mediated actin polymerization activity activated by VCA domains diffusing in the cytoplasm or bulk solution can be effectively suppressed. To test this idea, we first used mammalian culture cells (Madin-Darby Canine Kidney; MDCK cells), which contain all the components necessary for actin polymerization. As an optogenetic switch, we employed an improved Light-Induced Dimer (iLID) system, in which iLID and SspB bind together upon blue light illumination and reversibly dissociate under dark conditions (Guntas et al. 2015).

**Figure 1.**
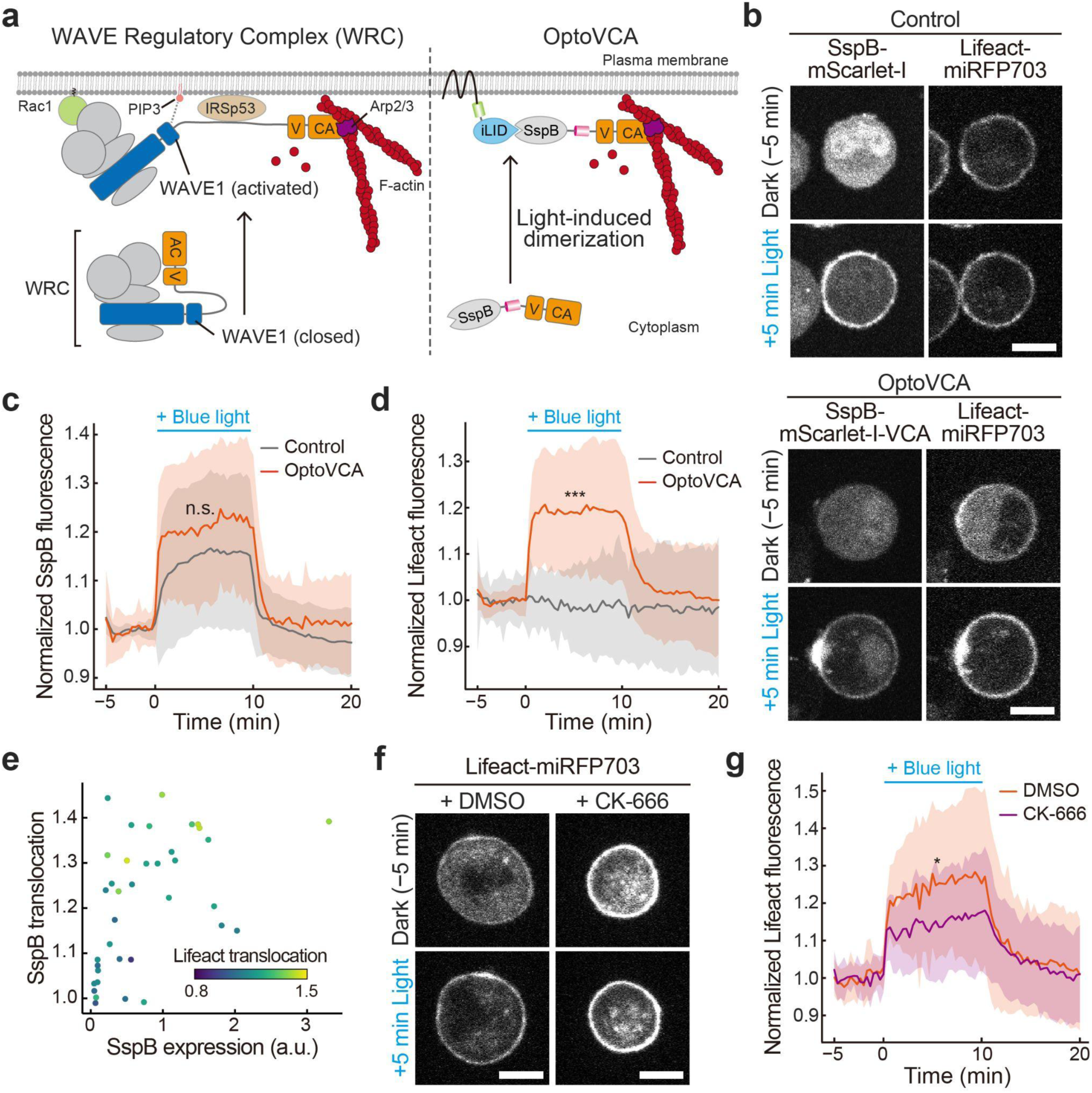
Light-induced actin polymerization in live cells. (a) Comparison of the regulatory mechanisms of WRC (left) and OptoVCA system (right). (b) Representative images of MDCK cells simultaneously expressing Stargazin-mEGFP-iLID and Lifeact-miRFP703 (right panels) along with SspB-mScarlet-I (upper left) or SspB-mScarlet-I-VCA (lower left). For each cell, protein localizations before (upper) and after (lower) blue light illumination are shown. (c, d) Normalized translocation levels of SspB-mScarlet-I (c, Control) or SspB-mScarlet-I-VCA (c, OptoVCA) and Lifeact-miRFP703 (d) to the cortex. The mean values (bold lines) are plotted as a function of time with the SD. *n* = 37 for each cell. (e) Relationship between the normalized translocation levels of SspB-mScarlet-I-VCA and Lifeact-miRFP703, and the expression level of SspB-mScarlet-I-VCA. The translocation levels of SspB and Lifeact are defined as the mean values of SspB-mScarlet-I-VCA and Lifeact-miRFP703 at *t* = 2-8 min in (c) and (d), respectively. *n* = 37 cells. (f) Representative images of the cells expressing Stargazin-mEGFP-iLID, SspB-mScarlet-I-VCA, and Lifeact-miRFP703 treated with DMSO (left) or CK-666 (right) before (upper) and after (lower) blue light illumination. (g) Normalized cortical fluorescence intensity of Lifeact-miRFP703 in (f). The mean values (bold lines) are plotted as a function of time with the SD. *n* = 18 and 17 for DMSO and CK-666 treated cells, respectively. All scale bars, 20 μm. *p* values were calculated by unpaired two-sided *t*-test at *t* = 5 min. ***, *p* < 0.001. *, *p* < 0.05. n.s., *p* ≥ 0.05.

To manipulate the intracellular localization of the VCA domain and to increase its local density on the plasma membrane, we expressed iLID fused with stargazin (Yamamoto et al. 2021; Natwick and Collins 2021) and SspB fused with the VCA domain (hereafter referred to as the OptoVCA; Fig. 1a, S1a). Each fusion protein was visualized using mEGFP or mScarlet-I, respectively. To evaluate the effect of the translocation of the VCA domain on the cortical F-actin dynamics, we used MDCK cells expressing Lifeact-miRFP703 and Stargazin-mEGFP-iLID immediately after the trypsinization. SspB-mScarlet-I-VCA or SspB-mScarlet-I were also co-expressed as OptoVCA and the control, respectively. As a result, both SspB-mScarlet-I-VCA and SspB-mScarlet-I translocated from the cytoplasm to the plasma membrane upon blue light illumination (Fig. 1b, 1c, S1b, S1c). On the other hand, the cortical fluorescence intensity of Lifeact-miRFP703 was increased only in the cells expressing SspB-mScarlet-I-VCA and reached a nearly steady state within ∼2 min (Fig. 1b, 1d, S1b, S1d). After stopping the illumination, both the SspB and Lifeact signals returned to the basal levels within ∼4 min. These results demonstrate that the OptoVCA system can rapidly and reversibly control actin polymerization and depolymerization.

In these experiments, we transiently expressed Stargazin-mEGFP-iLID and SspB-mScarlet-I-VCA. Consequently, the expression ratio of these optogenetic proteins and the expression level of SspB-mScarlet-I-VCA varied among the cells. To clarify the relationship between the expression level and the translocation efficiency of the SspB-mScarlet-I-VCA, and the resulting actin polymerization, we analyzed these parameters in each cell. The result shows that the cells with higher expression levels and translocation efficiency positively correlate with a higher increase in F-actin fluorescence (Fig. 1e). This further supports the significance of the local increase in the VCA density for efficient actin polymerization.

Finally, we confirmed that this actin polymerization is mediated by the Arp2/3 complex; the cells treated with CK-666, an Arp2/3 complex inhibitor, showed a relatively weaker increase in the cortical Lifeact signal compared to the DMSO-treated control cells during illumination (Fig. 1f, 1g), although the translocation dynamics of SspB-mScarlet-I-VCA were nearly identical between the two conditions (Fig. S1e, S1f). In addition, we frequently observed wavy shape changes in the cells expressing OptoVCA upon illumination (Fig. S1g). This result is consistent with the previous report showing that artificially localized VCA domain to the actin cortex softens and deforms mouse oocytes (Chaigne et al. 2015). Collectively, these results show that the OptoVCA system can control actin polymerization by recruiting the Arp2/3 complex to beneath the plasma membrane with blue light.

### Optogenetic manipulation of actin polymerization on a supported lipid bilayer

Next, we expanded this optogenetic approach to an *in vitro* reconstitution system. We combined the OptoVCA with a supported lipid bilayer (SLB), and reconstructed a biologically relevant actin network from purified proteins on the lipid membrane. As for the SLB, we used POPC as the main component and added 2-6% of DGS-NTA(Ni) to anchor His-mEGFP-iLID on the SLB (Fig. S2a). The smoothness of the SLB was confirmed by the fluorescence signals of Marina Blue-conjugated phospholipids and His-mEGFP-iLID attached to the membrane after forming the SLB (Fig. S2b). The fluidity of the SLB was also confirmed by observing the lateral Brownian motion of single His-mEGFP-iLID molecules with low density along the membrane using TIRF microscopy (Fig. S2c). To quantify the diffusion coefficient of His-mEGFP-iLID on the SLB under the saturated density, we replaced mEGFP with ShadowG, a dark variant of mEGFP, and mixed the protein with His-mEGFP- iLID at a molecular ratio of 1000:1 (Murakoshi et al. 2015) (Fig. S1d, S1e). The obtained diffusion coefficient of His-mEGFP-iLID was approximately 0.38 μm^2^/sec, which is comparable with the diffusion coefficient of membrane-bound proteins (Ziemba and Falke 2013) (Fig. S1f, S1g). The density of His-mEGFP-iLID on the SLB was controllable by changing the percentage of DGS- NTA(Ni) in the SLB (Fig. S2h, S2i). We next examined if flavin mononucleotide (FMN) is needed as a chromophore for iLID (Guntas et al. 2015). We loaded SspB-mScarlet-I-VCA to the His- mEGFP-iLID bound on the SLB in the presence of various FMN concentrations, then we illuminated a 37 × 37 μm square pattern of blue light at each condition and quantified the fluorescence intensities of SspB-mScarlet-I-VCA recruited on the SLB. Notably, the addition of >1 μM FMN dramatically improved the translocation efficiency of SspB-mScarlet-VCA to the SLB (Fig. S2j, S2k, S2l). Consistently, the binding of added FMN to purified iLID was detected by measuring absorbance at 450 nm using UV-vis spectroscopy (Fig. S2m, S2n). Therefore, we added 10 μM FMN in the following experiments. Finally, we quantified the extent to which SspB-mScarlet-I-VCA diffused outside the illuminated region due to membrane fluidity. Line scans showed that the density of SspB- mScarlet-I-VCA gradually decreased from the edge of the illuminated region over a distance of 10- 20 μm (Fig. S2o, S2p). Within 5 minutes after stopping the illumination, most of the bound SspB- mScarlet-I-VCA dissociated from the SLB.

To induce actin network assembly by blue light illumination, we added actin, profilin, the Arp2/3 complex, and capping protein (CP) to the OptoVCA. The cytoskeletal protein components were chosen according to the previous reports (Boujemaa-Paterski et al. 2017; Bieling et al. 2016; Dürre et al. 2018) (Fig. 2a). Profilin binds to actin monomers and suppresses the spontaneous nucleation in bulk solution, while CP binds to the barbed ends of actin filaments and suppresses excessive filament elongation. Before conducting the reconstitution experiments, we measured the actin polymerization activity of the purified proteins by the pyrene fluorescence assay (Fig. S3a), and confirmed that the polymerization activity is not affected by the addition of FMN or His-mEGFP- iLID, but is dependent on the presence of SspB-mScarlet-I-VCA and the Arp2/3 complex. Upon blue light illumination, SspB-mScarlet-I-VCA was rapidly recruited to the lipid surface, with the density nearly proportional to the percentage of DGS-NTA(Ni) in the SLB (Fig. 2b, 2c, 2d). Subsequently, actin polymerization started in the illuminated region (Fig. 2b, 2c, 2e). Notably, actin polymerization on the 6% DGS-NTA(Ni) lipid initiated much faster than that on the 2% DGS-NTA(Ni) lipid; the visible polymerization started within 1 min (Fig. 2b, 2c, 2e). After starting the polymerization on the 6% DGS-NTA(Ni) lipid, the fluorescence intensity of actin increased transiently and then decreased near the bottom plane close to the SLB (Fig. 2e). This phenomenon can be attributed to the local protein depletion effect around the illuminated region; the higher the VCA density on the SLB, the more monomer actin and ABPs are recruited to the SLB, leading to a decrease in the surrounding protein concentrations (Boujemaa-Paterski et al. 2017; Manhart et al. 2019; Dimchev et al. 2017). On the other hand, the slower increase in actin density on the 2% DGS-NTA(Ni) lipid suggests that the depletion effect did not occur because actin molecules in the bulk solution were slowly consumed, eventually reaching a steady state (Fig. 2c, 2e). During illumination, the polymerized actin gradually formed a cup-like or dome-like structure on the 2% and 6% DGS-NTA(Ni) lipid, respectively (Fig. 2f). Without light illumination or in the absence of the Arp2/3 complex, actin fluorescence was nearly undetectable (Fig. S3b, S3c, S3d). This clarified that actin polymerization in the *in vitro* OptoVCA system is also mediated by the recruitment of SspB-mScarlet-I-VCA and Arp2/3 complex to the membrane.

**Figure 2.**
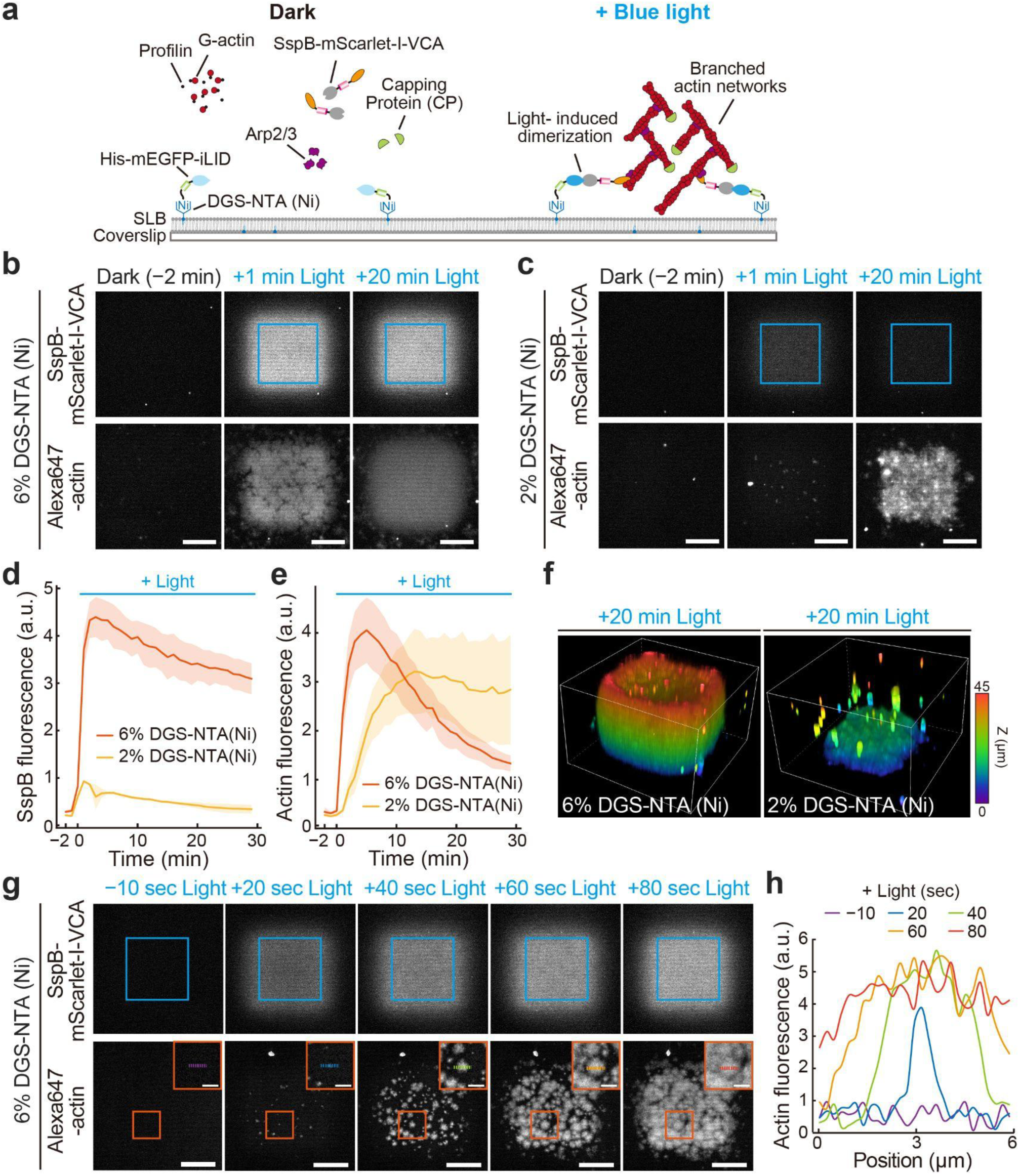
Light-induced actin polymerization on a supported lipid bilayer (SLB). (a) Schematic illustration of the OptoVCA system for *in vitro* experiments. Upon blue light illumination, actin polymerization is triggered by the recruitment of SspB-mScarlet-I-VCA to the SLB. Condition: 5 μM Actin, 15 μM Profilin, 100 nM Arp2/3 complex, 25 nM CP, and 150 nM SspB-mScarlet-I-VCA. (b, c) Representative images of the recruitment of SspB-mScarlet-I-VCA (upper panels) and subsequent actin polymerization (lower panels) on 6% DGS-NTA(Ni) lipid (b) and 2% DGS-NTA(Ni) lipid (c) at each indicated time point. Blue squares indicate the light-illuminated regions with 1 nW/μm^2^. All scale bars, 20 μm. (d, e) Fluorescence intensities of SspB-mScarlet-I-VCA (d) and Alexa647-actin (e) on 2% or 6% DGS-NTA(Ni) lipid. The mean values (bold lines) are plotted as a function of time with the SD. *n* = 12 for each condition. (f) Three-dimensional reconstructed image of polymerized actin on 6% DGS-NTA(Ni) lipid (left) and 2% DGS-NTA(Ni) lipid (right) after 20 min light illumination. (g) High-speed imaging of the recruitment of SspB-mScarlet-I-VCA (upper panels) and subsequent actin polymerization (lower panels) on 6% DGS-NTA(Ni) lipid. Blue-squared regions were illuminated with 1 nW/μm^2^. Orange-squared regions in the lower panels are magnified and shown as the inset images. Scale bars, 20 μm and 5 μm for the wide-view images and the inset images, respectively. (h) Growth dynamics of a single actin network seed quantified by the line scan along red dashed lines in the inset images of (g).

Finally, to dissect the initiation process of the actin network assembly, we observed the growth of polymerized actin with high-speed imaging. We found that actin clusters appeared on the SLB within 20 sec, then gradually expanded and fused with each other (Fig. 2g, 2h, Movie S1). Considering that the nucleation process requires a mother actin filament, the Arp2/3 complex, and the VCA domain, this observation suggests that the nucleation is initiated by the binding of diffusing F-actin fragments stochastically nucleated in the bulk region to the complex of the Arp2/3 complex and two VCA domains formed on the SLB (Gautreau et al. 2022; Espinoza-Sanchez et al. 2018). Collectively, these results demonstrate that the optimized setup of the OptoVCA system enables flexible control of actin polymerization in both spatial and temporal manners *in vitro*.

### Temporal control of the actin network assembly

To characterize the temporal resolution of the *in vitro* OptoVCA system, we examined how quickly actin polymerization halts after stopping the illumination. We illuminated the 6% DGS-NTA(Ni) lipid for 10, 20, or 30 min with a 37 × 37 μm square pattern (Fig. S4a). After stopping the illumination, the fluorescence intensity of SspB-mScarlet-I-VCA rapidly decreased (Fig. 3a), while the fluorescence intensity of actin gradually decreased depending on the illumination duration (Fig. 3b, 3d, S4b, S4c). This may be caused by the collapse of the actin network, such as debranching of the Arp2/3 complex-mediated branches (Le Clainche et al. 2003). As time progresses after the formation of the actin network and ATP hydrolysis proceeds within the filaments, the fraction of ADP-actin, known to have reduced affinity for the Arp2/3 complex, increases in the network (Blanchoin et al. 2000), resulting in debranching. We next focused on the growth dynamics of the actin pillar. While actin pillars continued to grow under constant light illumination, the growth ceased immediately after stopping the illumination (Fig. S4b, 3e, Movie S3). By changing the illumination pattern over time, the more complex actin structures were also freely designed (Fig. S4d, S4e, Movie S2). In this way, the OptoVCA system allows us to precisely design actin network thickness and three-dimensional structures.

**Figure 3.**
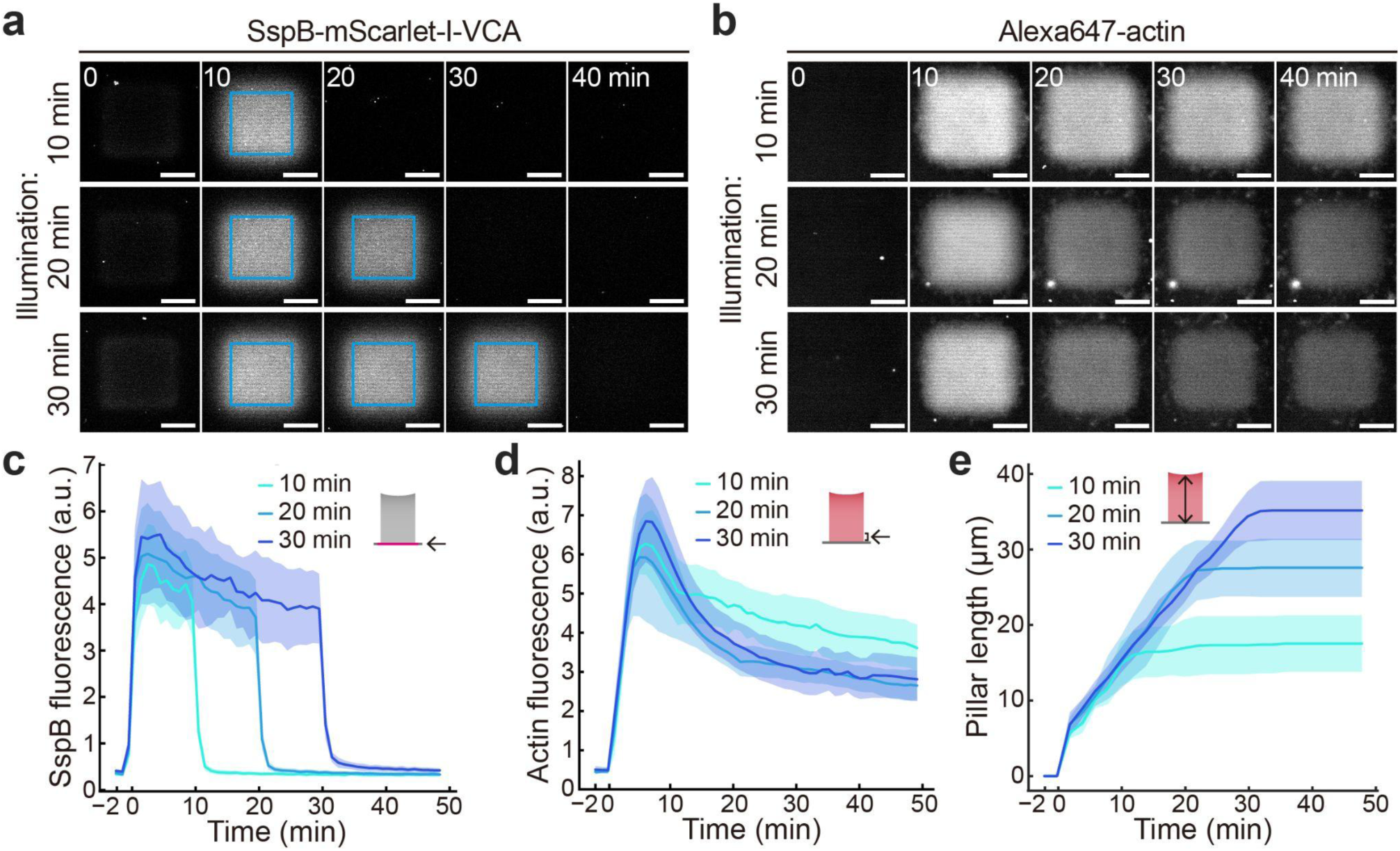
Temporal control of actin network assembly. (a, b) Representative images of the recruitment of SspB-mScarlet-I-VCA (a) and subsequent actin polymerization (b) at each indicated time point in three different illumination duration patterns. Blue squares indicate the light-illuminated regions with 1 nW/μm^2^. All scale bars, 20 μm. (c, d, e) Fluorescence intensities of SspB-mScarlet-I-VCA (c) and Alexa647-actin (d), and the pillar length (e) in three different illumination duration patterns. The mean values (bold lines) are plotted as a function of time with the SD. *n* = 12 for each condition.

### Control of the actin network density by the light intensity

To quantify the relationship between the local density of the VCA domain on the SLB and the resulting density of the actin network, we next examined whether our system can control the efficiency of SspB-mScarlet-I-VCA recruitment by varying the light intensity. Given the high sensitivity of the LOV domain used as a light-responsive protein in iLID (Berlew et al. 2022), we reduced the light intensity to ∼1% by inserting an ND filter into the light path of our microscope. We confirmed that the exact light intensity emitted from the objective lens was linearly associated with the output power controlled by the microscope software (Fig. S5a, S5b). By illuminating 25 × 25 μm square patterns of blue light with varied intensities to the 6% DGS-NTA(Ni) lipid, we found that the efficiency of SspB-mScarlet-I-VCA recruitment was almost saturated at the intensity with 0.15 nW/μm^2^ (Fig. 4a, 4b, 4c, Movie S4). On the other hand, we observed reduced recruitment of SspB-mScarlet-I-VCA at outputs of less than 0.15 nW/μm^2^. Consistent with this, the densities of polymerized actin were nearly proportional to the densities of SspB-mScarlet-I-VCA on the SLB when the outputs were smaller than 0.15 nW/μm^2^ (Fig. 4d, 4e). The fluorescence intensity of actin pillars assembled with an output greater than 0.11 nW/μm^2^ started to decrease 10 min after starting the illumination, presumably due to the local protein depletion effect (Fig. 4d). A similar trend was observed when a single square pattern of the same size was illuminated at the center of the microscopic field of view, supporting that the depletion effect occurs independently of the number and size of illuminated patterns (Fig. S5c, S5d). The growth rate of actin pillars formed under the strong illumination was slower than that formed under the weak illumination (Fig. 4a, S5e). This result can be explained by the balance between the local VCA density on the SLB and the supply of actin monomers and ABPs from outside the pillars; if it is assumed that the supply of actin monomers and ABPs is homogenous around the pillars, the higher local VCA density consumes more actin monomers and ABPs, resulting in slower growth. Consistently, the total actin fluorescence intensity in the pillar cross-section remained comparable under illumination conditions of 0.11 nW/μm^2^ or higher (Fig. S5f). In addition, we noticed that the top of the pillars showed curved edges when illuminated with 0.15-0.18 nW/μm^2^ or without using the ND filter (Fig. 2f, 4a). As the illumination area decreased, the top surface of the actin pillar became flatter even under the highest illumination intensity (Fig. S5c). These results suggest that the depletion effect is more pronounced under conditions of higher light intensity and wider illumination area, and that the stronger depletion at the central region contributes to the formation of the curved edge. Leakage of SspB-mScarlet-I-VCA outside the illuminated region due to lateral diffusion may also contribute to the formation of the curved edge. Furthermore, we evaluated the effects of imaging intervals and the phototoxicity induced by blue light illumination on actin molecules. In the absence of the antioxidant Trolox, the growth of actin filaments was likely inhibited by phototoxicity, whereas in the presence of Trolox, actin fluorescence and growth remained similar regardless of the imaging interval (Fig. S5g). Therefore, phototoxicity affecting the actin network caused by the fluorescence imaging is negligible under our observation conditions. Overall, our system can precisely manipulate the actin network density by controlling the recruitment of SspB-mScarlet-I-VCA to the SLB through variations in light intensity.

**Figure 4.**
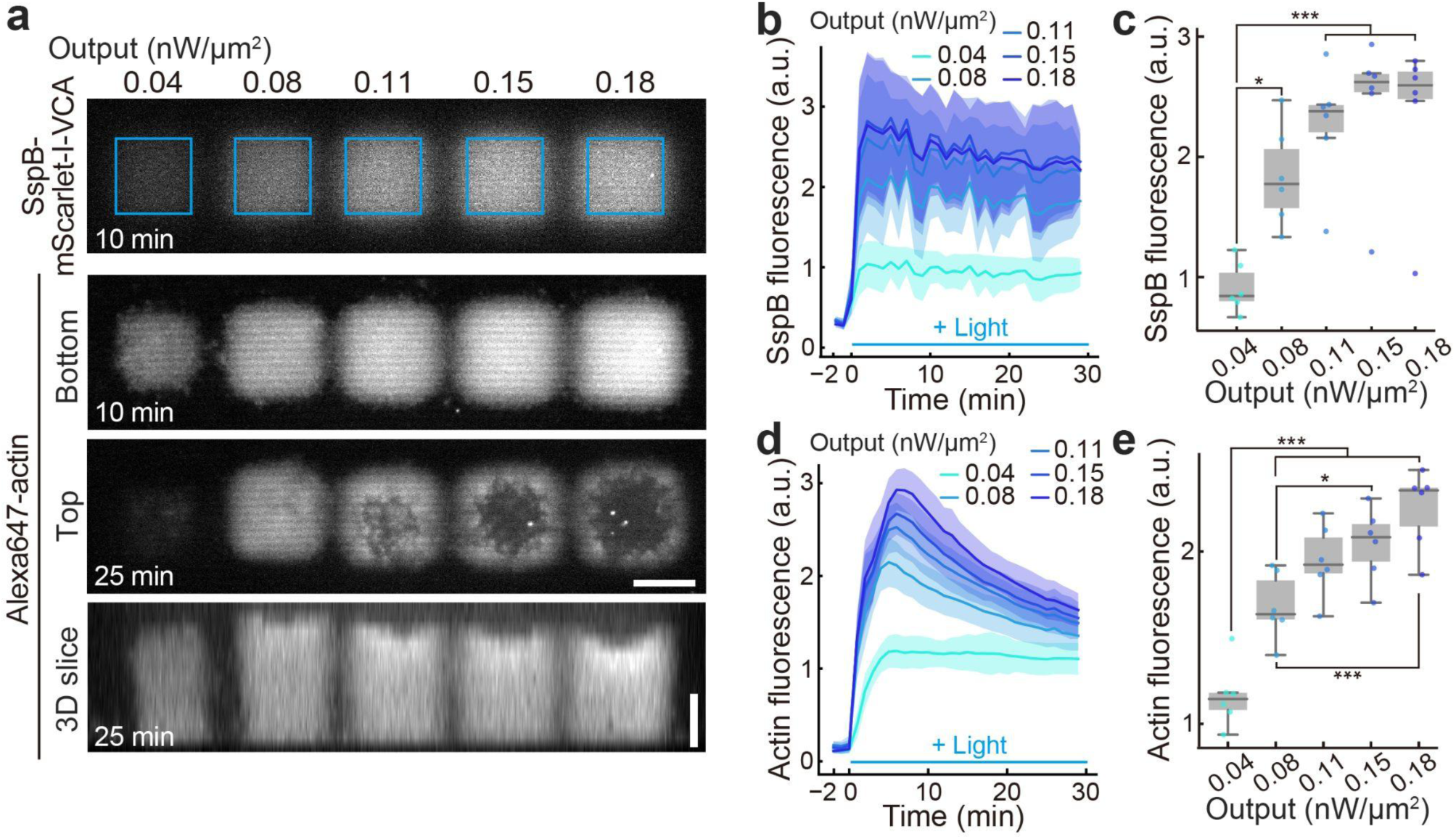
Control of actin density by light intensity. (a) Representative images of the recruitment of SspB-mScarlet-I-VCA (upper panel) and subsequent actin polymerization (lower panels) under five different light-powered illuminations on 6% DGS-NTA(Ni) lipid. The bottom plane, top plane, and three-dimensional(3D) reconstructed slice images at each indicated time point are shown. Condition: 5 μM Actin, 15 μM Profilin, 100 nM Arp2/3 complex, 25 nM CP, and 150 nM SspB-mScarlet-I-VCA. Blue squares indicate the illuminated regions. All scale bars, 20 μm. (b, c) Fluorescence intensities of SspB-mScarlet-I-VCA under the five different light-powered illuminations. The extreme outliers in (c) are attributed to focus drift of the microscope. (d, e) Fluorescence intensities of Alexa647-actin with five different light-powered illuminations. (b, d) The mean values (bold lines) are plotted as a function of time with the SD. *n* = 6 for each condition. (c, e) For box plots in (c) and (e), the mean fluorescence intensities of SspB-mScarlet-I-VCA in (b) and Alexa647-actin in (d) at *t* = 11-20 min are used, respectively. *p* values were calculated by one-way ANOVA followed by Tukey’s multiple comparisons test. ***, *p* < 0.001. **, *p* < 0.01. *, *p* < 0.05.

### Actin density-dependent penetration of myosin filaments into the network

Next, we applied the OptoVCA system to understand the relationship between the density of the actin network, the size of ABPs, and their penetration ability into the network. In living cells, it has been suggested that the penetration of myosin filaments into the cortical actin network is crucial for generating proper contractile force (Truong Quang et al. 2021). Therefore, we first chose myosin II as a representative ABP and examined to what extent the actin network density impacts the myosin penetration ability and the network deformability. Myosin molecules assemble into filaments through the phosphorylation of their light chains (Craig and Woodhead 2006). Prior to the experiments, smooth muscle myosin (SMM) was phosphorylated by ZIP kinase to induce filament formation and activation of motor activity (Komatsu and Ikebe 2004; Murata-Hori et al. 1999). The length of myosin filaments was 1.32 ± 0.66 μm (mean ± SD; Fig. S6a, S6b). To examine the effects of actin network density on myosin distributions, we mixed the preformed myosin filaments with the other components, and then induced the formation of actin networks with various densities. To minimize the depletion effect, illumination was limited to 10 min, as the onset of depletion, which was indicated by a decrease in actin fluorescence intensity, was observed after ∼10 min of illumination (Fig. 4d). Line profiles of actin fluorescence intensity at the middle height of the actin pillar indicate that local depletion is negligible in the interior region (Fig. S6c, S6d, S6e). Interestingly, while the myosin signal was detected within the sparse actin network assembled with 0.04 nW/μm^2^ output, most of the myosin signal was detected on the surface of the dense actin networks assembled with 0.08 nW/μm^2^ and 0.15 nW/μm^2^ outputs (Fig. 5a, middle column, S6f). This result indicates that the actin network density determines the efficiency of myosin filament penetration into the network. Strikingly, only a ∼2.5-fold increase of the actin network density almost completely inhibits penetration of myosin filaments in an all-or-none fashion (Fig. 5a, middle column, Fig. 5b). To further explore the network density effect, we formed a sparser network by reducing the concentrations of actin and ABPs in the system (Fig. 5a, right column, 5b). As expected, in the sparse condition, the myosin signal was detected in the network even under illumination with 0.08 nW/μm^2^ and 0.15 nW/μm^2^ outputs. Notably, the myosin and actin signals in the sparse conditions were mutually exclusive (Fig. 5c). Next, we asked whether the penetration ability of myosin filaments is dependent on only the actin network density or also on the ATPase activity of myosin motors. To examine this, we treated myosin filaments with blebbistatin, a myosin ATPase inhibitor (Straight et al. 2003), and compared the myosin signal intensity inside the actin networks with control conditions (Fig. S6g). In the networks assembled with 0.08 nW/μm^2^ and 0.15 nW/μm^2^ outputs, the myosin signal was significantly weaker in the blebbistatin-treated condition than in the control condition (Fig. S6h, S6i). These results provide direct evidence that both a sparse actin network and myosin with ATPase activity contribute to the efficient penetration of myosin filaments into the actin network.

**Figure 5.**
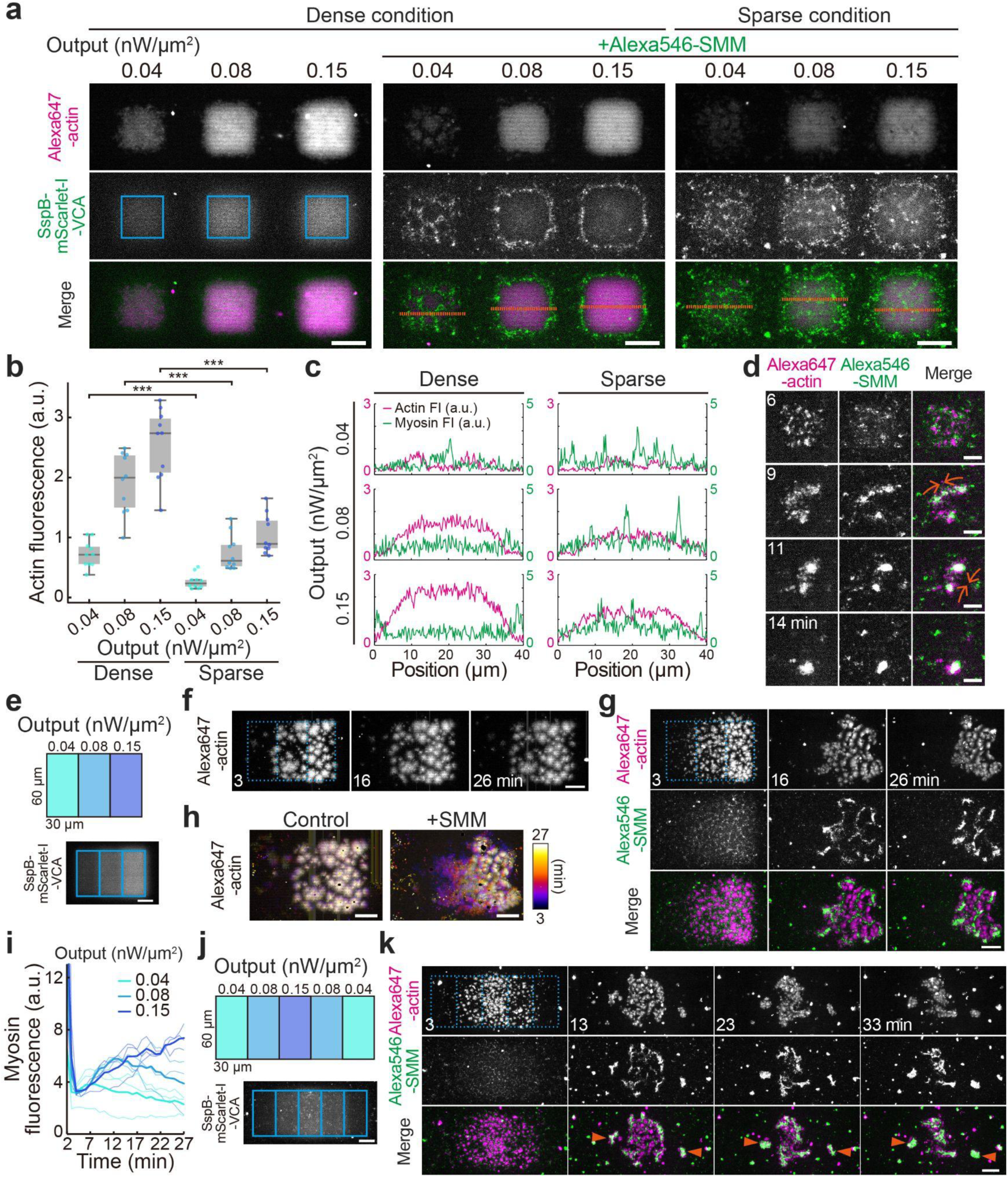
Actin density-dependent myosin penetrations and flows. (a) Representative images of polymerized actin (upper panels) and SspB-mScarlet-I-VCA (middle panels) in the absence of (left column) or the presence of 200 nM Alexa546-SMM (middle and right columns). Dense condition: 5 μM Actin, 15 μM Profilin, 100 nM Arp2/3 complex, 25 nM CP, and 150 nM SspB-mScarlet-I-VCA. Sparse condition: 2 μM Actin, 6 μM Profilin, 100 nM Arp2/3 complex, 10 nM CP, and 150 nM SspB-mScarlet-I-VCA. Images taken after 5 min illumination are shown. Blue squares indicate the illuminated regions. (b) Box plots of the fluorescence intensities of actin 2 μm above SLB after 5 min illumination. *n* = 10 for each condition. *p* values were calculated by unpaired two-sided *t*-test. ***, *p* < 0.001. (c) Line profiles of the actin and myosin intensities along the orange dashed lines shown in (a). (d) Representative images of the movements of actin and myosin puncta. Blue light was illuminated for 5 min from *t* = 0 to 5 min. The elapsed times are indicated in the images of Alexa647-actin. Orange arrows indicate the direction of the puncta movements. (e) Illumination pattern to induce actin flow (upper) and the fluorescence image of recruited SspB-mscarlet-I-VCA (lower). (f, g) Maximum intensity projection images of the actin structure produced by the illumination pattern shown in (e) in the absence (f) or the presence of 200 nM Alexa546-SMM (g). The SMM signal is shown in middle panels in (g). The protein concentrations are identical to the sparse network in (a). Blue light was illuminated for 2 min from *t* = 0 to 2 min. The illuminated region was indicated as blue dashed boxes. The elapsed times are indicated in the images of Alexa647-actin. (h) Temporal color-coded images of the actin structure in (f, g) after stopping the illumination. (i) Fluorescence intensities of Alexa546-SMM in the three regions shown in (e) after stopping the illumination. The bold lines indicate the mean values of three independent experiments (thin lines). (j) Illumination pattern to induce actin flow from both sides (upper) and the fluorescence image of recruited SspB-mscarlet-I-VCA (lower). (k) Maximum intensity projection images of the actin and SMM structures produced by the illumination pattern shown in (j). Blue light was illuminated for 2 min from *t* = 0 to 2 min. The illuminated region was indicated as blue dashed boxes. The elapsed times are indicated in the images of Alexa647-actin. Orange arrowheads indicate disconnected actin structures in the sparse regions. Scale bars, 20 μm and 10 μm for (a, e, f, g, h, j, k) and (d), respectively.

Next, we evaluated how the network density impacts the deformability. Interestingly, after stopping the illumination with 0.04 nW/μm^2^ output in the sparse condition, the actin and myosin puncta were rapidly concentrated and formed a large cluster (Fig. 5d). This is consistent with the previous observations regarding the formation of actomyosin condensates in *in vitro* systems and living cells (Soares e Silva et al. 2011; Kruse et al. 2024). Such condensates were not formed in the dense networks, suggesting that the network deformability is determined by the penetration ability of myosin filaments. In living cells, myosin-driven network deformations are frequently observed as a cortical flow during migration and cytokinesis (Kelkar, Bohec, and Charras 2020; Paluch, Aspalter, and Sixt 2016). However, the causal relationship between the actin network density and the direction of the cortical flow has not been explicitly demonstrated due to the complexity of living cells. To examine the direct role of the network density in the actin flow, we assembled the actin network having a density gradient by changing the illumination power in space (Fig. 5e). To obtain a thin deformable actin sheet, we illuminated the SLB for 1-2 min. In the absence of myosin, we confirmed that the actin sheet remained at the same position on the SLB for at least 24 min after stopping the illumination (Fig. 5f). In the presence of myosin, actin and myosin started contraction and concentrated toward the densest network region immediately after stopping the illumination (Fig. 5g, 5h, S6j, Movie S5). The fluorescence intensity of myosin decreased over time in the regions illuminated with 0.04 nW/μm^2^ and 0.08 nW/μm^2^ outputs, whereas it increased in the region illuminated with 0.15 nW/μm^2^ output, indicating that myosin filaments were transported toward the densest network region by unidirectional actin flow (Fig. 5i). To further demonstrate that the flow direction can be determined solely by the network density gradient, we illuminated strong light in the center region of the SLB and formed actin density gradients along both sides (Fig. 5j). As a result, the actin sheet showed flows toward the center, further supporting that the actin density gradient determines the flow direction (Fig. 5k, Movie S6). Moreover, we found that F-actin in the sparsest regions on both sides was not connected to the network and was freely floating (Fig. 5k, Movie S6). These results provide direct evidence that only the actin density gradient and the network connectivity are sufficient factors to induce the directed actin flow by myosin.

### Actin density-dependent disassembly of the network by cofilin

To examine the effect of relatively smaller-sized ABPs on the actin network, next we focused on cofilin, an actin depolymerization factor, with ∼3.5 nm in diameter (Tanaka et al. 2018; Huehn et al. 2020). We illuminated the SLB with five different outputs to assemble actin networks with different densities, and then monitored the network disassembly dynamics after stopping the illuminations both in dense and sparse conditions (Fig. 6a, 6b). For simplicity, blue light was illuminated only for 10 min to avoid the local protein depletion effect. Line profiles of actin fluorescence intensity at the middle height of the actin pillar suggests that local depletion is negligible inside the pillar (Fig. S7a, S7b, S7c). We quantified the actin network disassembly level in various network densities by calculating the ratio of the fluorescence intensities of actin before and 10 min after stopping the illuminations. Interestingly, the network disassembly level sharply decreased as the network density increased, showing that the cofilin-mediated network disassembly dramatically suppressed in dense actin networks (Fig. 6c). In addition, we found that the actin density just after stopping the illumination was nearly identical regardless of cofilin concentrations in the dense condition, while it was decreased with increasing cofilin concentrations in the sparse condition (Fig. S7d, S7e). These results are consistent with the previous report that the network disassembly is inversely proportional to the square of the local actin density and proportional to the actin-bound cofilin density (Manhart et al. 2019).

**Figure 6.**
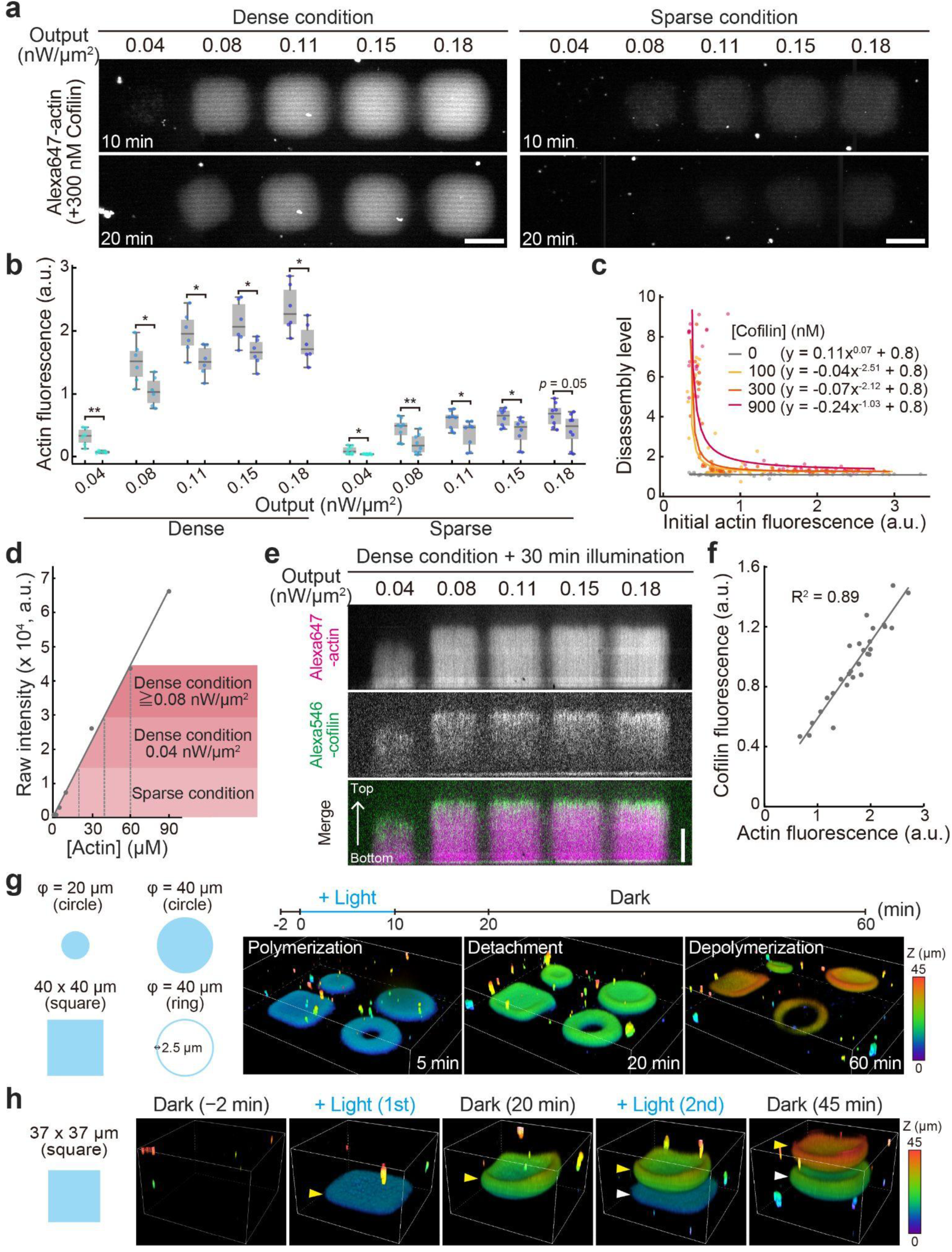
Effects of cofilin on various actin densities. (a) Maximum intensity projection images of actin networks after 10 min illumination (upper panels) and 10 min after stopping the illumination (lower panels). Dense condition: 5 μM Actin, 15 μM Profilin, 50 nM Arp2/3 complex, 25 nM CP, 150 nM SspB-mScarlet-I-VCA, and 300 nM Cofilin. Sparse condition: 2 μM Actin, 6 μM Profilin, 50 nM Arp2/3 complex, 10 nM CP, 150 nM SspB-mScarlet-I-VCA, and 300 nM Cofilin. All scale bars, 20 μm. (b) Box plots of the fluorescence intensities of Alexa647-actin after 10 min illumination (left boxes) and 10 min after stopping the illumination (right boxes) in the presence of 300 nM cofilin. *n* = 6 and 8 for dense and sparse conditions, respectively. *p* values were calculated by unpaired two-sided *t*-test. **, *p* < 0.01. *, *p* < 0.05. (c) Relationship between the initial fluorescence intensity of Alexa647-actin and the actin disassembly level. The disassembly level was calculated by dividing the intensity 10 min after illumination by the intensity 10 min after stopping the illumination. Bold lines indicate power-law fitting. *n* = 51, 60, 57, 42 for 0, 100, 300, 900 nM Cofilin, respectively. (d) Calibration curve to estimate the actin network density. Each color corresponds to the range of actin density in indicated illumination powers. (e) Spatial distribution of cofilin in the dense condition. Condition: 5 μM Actin, 15 μM Profilin, 100 nM Arp2/3 complex, 25 nM CP, 150 nM SspB-mScarlet-I-VCA, and 900 nM Alexa546-cofilin. Scale bar, 20 μm. (f) Relationship between the fluorescence intensities of actin and cofilin in single actin networks. *n* = 28 from individual actin networks. (g) Three-dimensional reconstructed images of floating actin networks on 6% DGS-NTA(Ni) lipid at each indicated time point produced by various illumination patterns. (h) Three-dimensional reconstructed images of floating actin networks by a square-shaped illumination pattern on 6% DGS-NTA(Ni) lipid at each indicated time point. Yellow and white arrowheads indicate the first and second polymerization, respectively. (g, h) Condition: 5 μM Actin, 15 μM Profilin, 100 nM Arp2/3 complex, 25 nM CP, 150 nM SspB-mScarlet-I-VCA, and 600 nM Cofilin.

Why is the cofilin-mediated network disassembly suppressed in dense actin networks? We hypothesized that the accessibility of cofilin to the actin pillar is sterically hindered by the network density at a specific threshold, as seen in myosin filaments. To test this hypothesis, we first estimated the actin network density by comparing the fluorescence intensities of actin in the pillars and homogeneous F-actin solutions with known concentrations under the same microscope (Fig. 6d, S8a). The estimated actin density in the dense network is ∼50 μM. This means that actin filaments are spaced ∼110 nm apart on average (Supplementary Note). Considering that the diameter of actin filaments is ∼8 nm (Blanchoin et al. 2014), this estimated value would be sparse enough for cofilin to penetrate into the network by diffusion. As expected, cofilin labeled with a small fluorescence dye penetrated into the dense network (Fig. 6e). Indeed, the cofilin density was proportional to the actin density in a wide range from sparse to dense networks, indicating that cofilin can freely diffuse into the network even in dense conditions (Fig. 6f). Notably, cofilin was enriched near the top of actin pillars. This may be because cofilin has a higher affinity to ADP-actin than ATP-actin (Carlier et al. 1997). Collectively, these results clarified that the actin network disassembly mediated by cofilin is also regulated by the network density. However, in the case of tiny cofilin, not the steric hindrance, but the network structure such as the branching frequency or the degree of network entanglement might contribute to the density-dependent suppression of the network disassembly.

Finally, to investigate whether the resistance to the cofilin-mediated network disassembly in the dense network is influenced by the network shapes, we illuminated the SLB with various patterns. Interestingly, the assembled network gradually floated, presumably due to actin disassembly preferentially from the network surface, and kept their shapes in any patterns for over 40 min (Fig. 6g, Movie S7). Although the density of ADP-actin is expected to be low at the bottom plane of the actin network, the floating behavior can be at least partially explained by a mechanism where diffusing cofilin depolymerizes actin at the bottom surface of the network, thereby detaching the network from the SLB. Moreover, the floating actin networks were repeatedly created from the same position by on/off cycles of the illumination (Fig. 6h, S7f, Movie S8). These results indicate that the effect of cofilin on the network disassembly is dependent on the network density but independent of the network shapes.

## Discussion

In this study, we developed an optogenetic tool, OptoVCA, which enables us to manipulate the assembly of the Arp2/3 complex-mediated actin networks by illuminating blue light. The strategy is applicable for both mammalian culture cells and *in vitro* reconstitution systems composed of purified actin cytoskeletal proteins and the SLB. We showed that the OptoVCA system can precisely control the density, thickness, and shape of actin networks on the same SLB by changing the illumination powers, durations, and patterns, respectively. Several papers have reported the techniques to localize NPFs to specified areas on the SLB or the glass surface, and succeeded in forming the Arp2/3 complex-mediated actin networks *in vitro* (Yamazaki et al. 2024; Colin et al. 2023; Boujemaa-Paterski et al. 2017; Bieling et al. 2016; Manhart et al. 2019; Anne-Cécile Reymann et al. 2010). However, the removal of NPFs from the substrate was technically limited, making it difficult to stop actin polymerization during the observation. Our optogenetic approach has overcome this limitation by simply stopping the illumination, enabling us to create the actin network with a desired thickness. Therefore, a thin actin structure such as the actin cortex can be reconstituted by our approach (Fig. 5e-k). Furthermore, the OptoVCA system enables the formation of actin structures with two-dimensional (Fig. 5f, 5g, 5k) or three-dimensional density gradients by spatially and temporally controlling light intensity. Furthermore, by dynamically changing the illumination area during imaging, more complex three-dimensional actin structures can be generated (Fig. S4d). More biologically relevant actin networks can also be reconstituted by combining other NPFs and crosslinker proteins with the OptoVCA. However, there are two remaining issues to be addressed in the OptoVCA system. First, in our current design, the excess SspB-mScarlet-I-VCA is diffusing in the bulk region so that actin polymerization in the bulk region cannot be completely suppressed (Fig. S3b). Bright aggregations frequently observed in the bulk region may be caused by such a mechanism and it may reduce the effective concentrations of the loaded proteins (Fig. 2f, 6g, 6h, S3b, S3c, S4a, S4d). Consistently, the overexpression of the SspB-mScarlet-I-VCA in mammalian cells enhanced cytoplasmic signals of the Lifeact-miRFP703 under dark conditions (Fig. 1b). Further improvement of the protein design, such as the light-induced formation of functional VCA domain through the dimerization of split V domain and CA domain, is needed to suppress the basal activity of the VCA domain. Second, in *in vitro* experiments, lateral diffusion of the recruited VCA domain from the illuminated regions leads to a spatial gradient of the recruited VCA domain and actin density at the periphery of the illuminated regions (Fig. S2p, S6d, S6e, S7b, S7c). This gradient arises from a technical limitation inherent to the use of a fluid lipid membrane. Incorporating the spatially patterned SLB into the OptoVCA would overcome this limitation (Yamazaki et al. 2024).

Using the OptoVCA system, we examined whether myosin filaments and cofilin are excluded from the actin network depending on the network density (Fig. 5, 6). The maximum actin density in our experiments was estimated to be ∼50 μM (Fig. S8a). The estimated density is relatively low compared to the F-actin concentration in lamellipodium ∼500 μM (Pollard 2016; Abraham et al. 1999; Koestler et al. 2009). Since actin network density strongly depended on the VCA density on the SLB (Fig. 2e, 4e), we also estimated the maximum density of SspB-mScarlet-I-VCA. While the iLID intensity increased proportional to the density of DGS-NTA(Ni), the SspB intensity saturated on the SLB containing DGS-NTA(Ni) greater than 5%, suggesting that steric hindrance occurred between the proteins on the 5% DGS-NTA(Ni) (Fig. S8b, S8c, S8d). Assuming that the area covered with a single lipid molecule is 0.64 nm^2^ (König, Dietrich, and Klose 1997) and iLID is fully activated on the SLB, the saturated density of SspB-mScarlet-I-VCA on the 5% DGS-NTA(Ni) lipid is estimated to be ∼75,000 molecules/μm^2^. This density is approximately 2.5-fold higher than that measured in live cells (Arasada and Pollard 2011; Ditlev et al. 2012). Therefore, the possible reason for the relatively low actin network density may not be because of the lower density of VCA on the SLB but because of the other factors, such as lower concentrations of the Arp2/3 complex and F-actin fragments and the lack of volume exclusion effects from the cytoplasm. Even in such conditions, we observed that myosin filaments with the mean length of 1.32 μm were excluded and trapped at the side surface of the network in which actin filaments are spaced 110 nm apart on average (Fig. 5, S8a, Supplementary Note). On the other hand, under conditions where the actin filament spacing is estimated to be 160 nm or larger, we observed the localization of myosin filaments within the network (Fig. 5a, S8a, Supplementary Note). The threshold spacing distance determining whether myosin filaments can penetrate into the network is expected to be between 110 nm and 160 nm. This value is comparable to the minimum size of myosin filaments used in the experiments (Fig. S6b). Thus, our *in vitro* experiments provided direct evidence that the ability of myosin penetration into the actin network is strictly regulated by the network density. Strikingly, we found that only a ∼2.5-fold increase in the network density, in other words, only a ∼1.5-fold increase in the mean actin mesh size, drastically changed the ability of myosin penetration (Fig. 5a, 5b, 5c). This strengthens the idea that the exclusion of myosin observed in the cellular actin cortex is attributable to the steric hindrance effect (Truong Quang et al. 2021). Interestingly, the myosin signal was undetectable at the top surface of the pillar, while a strong myosin signal was detected at the side surface (Fig. S6f). This might be due to the difference in the orientation of actin filaments between the side surface and the top surface of the pillar. On the side surface, actin filaments form an organized structure in which filaments are partially aligned, allowing myosin filaments to remain stably attached. On the top surface, actin filaments may form a disorganized spiky structure, leading to rapid dissociation of bound myosin filaments (Fig. 7). This mechanism might also contribute to the exclusion of myosin filaments from the actin cortex in mouse oocytes observed when the overexpressed VCA domain was localized beneath the plasma membrane (Chaigne et al. 2015). Moreover, in sparse conditions, we found that the motor activity enhances the penetration ability of myosin filaments into the network (Fig. S6g, S6i). In this way, the penetration ability of myosin filaments is regulated by the actin network density, orientation of actin filaments, and myosin motor activity (Fig. 7).

**Figure 7.**
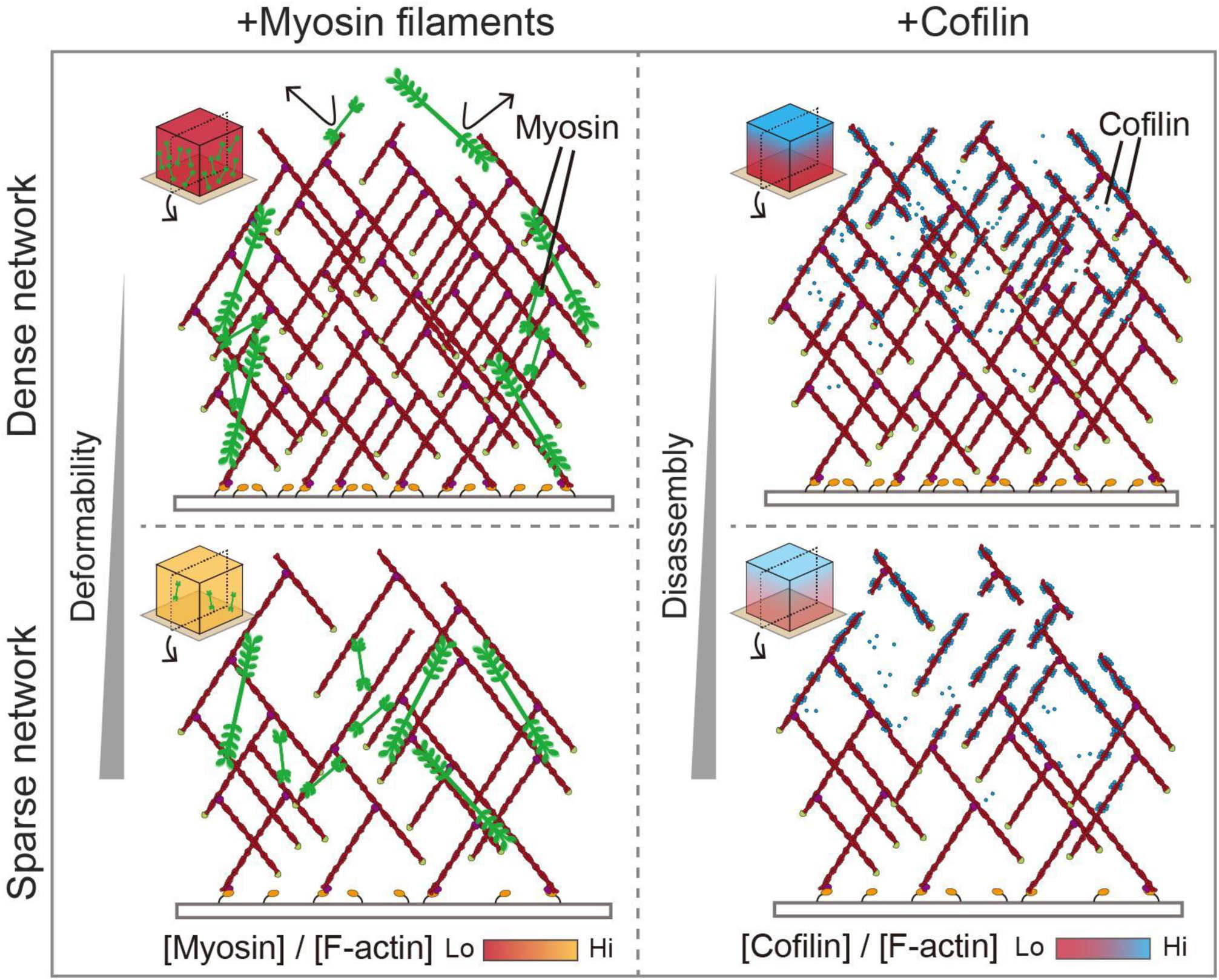
Actin network density-dependent protein penetrations and functions. Schematic illustration showing the overhead view and side view of assembled actin networks in the presence of myosin filaments (left) and cofilin (right). The color bars indicate the proportion of myosin and cofilin bound to actin filaments within the network. Myosin: In the dense network, myosin filaments cannot penetrate into the network and are trapped at the side surface due to the steric hindrance (upper left). Myosin filaments cannot stay longer on the top surface. In the sparse network, myosin filaments can penetrate into the network with the help of motor activity and the penetration makes the network deformable (lower left). Cofilin: cofilin can penetrate into both the sparse and the dense networks and preferentially binds to ADP-actin. In the dense network, the network disassembly rate is low (upper right). In the sparse network, cofilin-mediated network disassembly is more active (lower right).

In the case of cofilin, penetration was observed even in the densest condition (Fig. 6e). While the cofilin density was proportional to the actin density (Fig. 6f), the efficiency of the network disassembly was sharply decreased in the networks with the actin concentration above 20-30 μM. Therefore, this density-dependent suppression of cofilin-mediated network disassembly cannot be explained by the steric hindrance effect. To consider the possible mechanism, we estimate the filament level structure of the actin network used in the experiments. First, the mean length of actin filament branches in the dense condition is estimated to be ∼1-2 μm (Supplementary Note) (Pollard 2016; Schafer, Jennings, and Cooper 1996; Wear et al. 2003). Next, the density of actin molecules in the dense network is estimated to be ∼30,000/μm^3^. Therefore, the network is estimated to be composed of 40-80 actin filament branches per 1 μm^3^. There are two possibilities that can explain the density-dependent suppression mechanism of cofilin-mediated network disassembly. One possibility is that the severed actin fragments are less likely to escape from the dense network than from the sparse network because of the network entanglement if it is assumed that the frequency of the actin filaments severing by cofilin is the same between sparse and dense networks. Another possibility is the conformational difference in actin filaments between sparse and dense networks. The cofilin-mediated severing of actin filaments is known to be dependent on the twisting of actin (McGough et al. 1997). In dense networks, the higher frequency of the Arp2/3 complex-mediated branching of actin filaments and/or the entanglement with surrounding filaments may impair the twisting, potentially reducing the probability of cofilin-mediated severing of actin filaments. To further explore the network density-dependent effect of cofilin, it will be important to investigate the impact of ABPs on the severing activity of cofilin in single actin filament level and the diffusivity of F-actin fragments in the branched actin network.

We showed that the actin sheet flowed toward the denser network in the presence of myosin filaments (Fig. 5g, 5k). On the other hand, the sparsest network was disconnected from the directed flow (Fig. 5k). These observations clarified that controlling only two parameters, the network density gradient and the network connectivity, are sufficient to induce the directed flow and control the flow direction. In other words, our study clarified that the spatial regulation of myosin activity is not always required to induce the directed flow. In living cells, actin flow contributes to the deformation processes during migration, cytokinesis, and oogenesis (Munro, Nance, and Priess 2004; Wolke, Jezuit, and Priess 2007; Kelkar, Bohec, and Charras 2020; Paluch, Aspalter, and Sixt 2016). Such cellular deformability may be facilitated by regulating the spatial gradient of the actin network within a density range that myosin filaments can penetrate and the network is connected (i.e. dense condition with 0.04 nW/μm^2^ output or sparse condition with 0.04-0.15 nW/μm^2^ in Fig. 5a). In addition, as the actin flow proceeds, we observed the fusion and alignment of the myosin signal (Fig. 5g, 5k). It has been observed in mammalian cells that several myosin filaments stack into a single thick filament during the constriction of the contractile rings (Fenix et al. 2016; Yamamoto et al. 2019; Nilay et al. 2020). This stacked structure is considered to be important for the generation of sufficient contractile force (Fenix et al. 2016). The formation of the contractile ring is a sequential process that follows cortical flow from the poles to the equator within the actin cortex (Anne-Cecile Reymann et al. 2016; Bray and White 1988). The actin flow and myosin alignments observed in our study resemble this sequential process. Therefore, our study may highlight the importance of the density gradient of the actin network in the contractile ring formation process.

In summary, the OptoVCA system provides us with a flexible platform to reconstruct branched actin networks with various densities, thicknesses, and shapes. Taking these advantages, we provide direct evidence that the penetration ability of ABPs and their functions are tightly regulated by the actin network density. Moreover, by creating the actin sheet that has the density gradient, we identified the minimum factors sufficient to induce the directed actin flow. The actin network is broadly distributed throughout the cell with various densities, including the cortex, lamellipodia, and cytoplasm. Moreover, recent studies have increasingly suggested that the actin network regulates the distribution of organelles (Moore et al. 2021; Olguin-Olguin et al. 2021). Therefore, exploring the relationship between the network densities and protein or organelle sizes using an *in vitro* reconstitution system may help to understand not only the cell mechanics but also the compartmentalization mechanism within the cell in the future.

## Methods

### Plasmids

The oligonucleotides for PCR and DNA sequencing were purchased from Eurofin. The cDNAs of WAVE1 and cofilin were cloned from cDNA synthesized from mouse total RNA (Clontech). pCAGGS-Stargazin-mEGFP-iLID and pCAGGS-SspB-mScarlet-I were purchased from Addgene (#178523 and #178521, respectively). To construct pCAGGS-SspB-mScarlet-I-VCA, the VCA domain sequence of WAVE1 was inserted into pCAGGS-SspB-mScarlet-I using Gibson assembly cloning. For pET-28a(+)-His-mEGFP-iLID and pET-28a(+)-His-ShadowG-iLID, the cDNAs of mEGFP, ShadowG, and iLID were codon optimized for *E.coli,* synthesized (Eurofin), and inserted into the pET-28a(+) vector using Gibson assembly cloning. For pGEX-6P-1-SspB-mScarlet-I-VCA, the cDNAs of SspB and mScarlet-I were codon optimized for *E.coli* and synthesized (Eurofin), and inserted into the pGEX-6P-1 vector using Gibson assembly cloning. After that, the VCA domain sequence was amplified from pCAGGS-SspB-mScarlet-I-VCA and subsequently inserted into the vector using Gibson assembly cloning. To construct pRSET-mCofilin1, the cofilin sequence was inserted into the pRSET vector by Gibson assembly cloning. pGEX-5X-hZIPK was a kind gift from Dr. Kozue Hamao (Hiroshima University).

### Protein expression and purification

#### His-mEGFP-iLID and His-ShadowG-iLID

His-mEGFP-iLID and His-ShadowG-iLID were expressed in BL21 (DE3) in MMI medium. The cells were grown at 37°C until the OD600 was reached at 0.6. The protein expression was induced by the addition of 1 mM isopropyl-β-D-thiogalactopyranoside (IPTG). The cells were shaken at 160 rpm at 22°C for 20 hours. The collected cells were resuspended in PBS (pH 7.4) containing 20 mM imidazole-HCl pH 7.4, 10 μg/mL leupeptin, 10 μg/mL pepstatin, 10 μg/mL chymostatin, and 1 mM PMSF. After sonication and centrifugation, the supernatant was collected and loaded onto the HisTrap HP column (Cytiva), which was controlled by AKTA Start (Cytiva). After the column washes with 500 mM NaCl containing 20 mM imidazole-HCl pH 7.4, the protein was eluted with 500 mM NaCl containing 500 mM imidazole-HCl pH 7.4. The eluted protein fractions were concentrated using Amicon Ultra-4 (30 K MWCO, Millipore) and gel-filtrated over the Superdex 200 Increase 10/300 GL column with A50 buffer (50 mM HEPES-KOH pH7.6, 50 mM KCl, 5 mM MgCl_2_, 1 mM EGTA) containing 1 mM DTT, controlled by the AKTA Explorer. Finally, the protein was snap-frozen in liquid nitrogen and stored at –80°C.

#### SspB-mScarlet-I-VCA

SspB-mScarlet-I-VCA was expressed in BL21 (DE3) in MMI medium. The cells were grown at 37 °C until the OD600 was reached at 0.6. The expression was induced by the addition of 1 mM isopropyl-β-D-thiogalactopyranoside (IPTG). The cells were shaken at 160 rpm at 22 °C for 20 hours. The collected cells were resuspended in PBS (pH 7.4) containing 10 μg/mL leupeptin, 10 μg/mL pepstatin, 10 μg/mL chymostatin, and 1 mM PMSF. After sonication and centrifugation, the supernatant was collected and loaded onto the GSTrap HP column (Cytiva). After the column wash with PBS and 5 volumes of PreScission cleavage buffer (50 mM Tris-HCl pH7.5, 150 mM NaCl, 1 mM EDTA, 1 mM DTT), PreScission Protease (Cytiva) was injected into the column. Then, the column was incubated at 4 °C overnight to remove GST-tag. The cleaved protein was collected and gel-filtrated over the Superdex 200 Increase 10/300 GL column with A50 buffer containing 1 mM DTT, controlled by the AKTA Explorer. Finally, the protein was snap-frozen in liquid nitrogen and stored at –80 °C.

#### Actin

Actin was purified from the acetone-dried powder of rabbit skeletal muscle (M6890, Sigma) according to a previous report (Spudich and Watt 1971). The purified actin was labeled with Alexa Fluor 647 C5 Maleimide (Invitrogen). Labeled actin was clarified by gel-filtration over Superdex 200 Increase 10/300 GL column (Cytiva) in G-buffer (2 mM Tis-HCl pH8.0, 0.05 mM CaCl_2_, 2 mM NaN_3_, 0.1 mM ATP, 0.5 mM 2-mercaptoethanol). Both unlabeled and labeled actin in G-buffer were snap -frozen in liquid nitrogen and stored at –80 °C.

#### Cofilin

Cofilin was expressed in BL21 (DE3) in MMI medium. The cells were grown at 37 °C until the OD600 was reached at 1.0. The expression was induced by the addition of 1 mM IPTG. The cells were shaken with 130 rpm at 25 °C for 3 hours. The collected cells were resuspended in PBS (pH 7.4) containing 10 μg/mL leupeptin, 10 μg/mL pepstatin, 10 μg/mL chymostatin, and 1 mM PMSF. After sonication and centrifugation, the supernatant was collected and loaded onto the GSTrap HP column (Cytiva). After the column wash with PBS and 5 volumes of PreScission cleavage buffer, PreScission Protease (Cytiva) was injected into the column. Then, the column was incubated at 4 °C overnight to remove GST-tag. The cleaved protein was collected and concentrated using Amicon Ultra-4 (3 K MWCO, Millipore), and then gel-filtrated over the Superdex 200 Increase 10/300 GL column with A50 buffer containing 1 mM DTT, controlled by the AKTA Explorer. For fluorescence labeling, the protein was mixed with Alexa Fluor 546 NHS Ester (Invitrogen), and incubated at 4 °C overnight. The reaction was stopped by adding 10 mM Tris-HCl (pH 7.5) and the protein was gel filtrated at the same condition as the non-labeled protein. Both unlabeled and labeled cofilin were snap-frozen in liquid nitrogen and stored at –80 °C.

### Myosin

Unphosphorylated smooth muscle myosin (SMM) was purified from chicken gizzards as previously reported (Ikebe and Hartshorne 1985). The extracted sample was further dialyzed at 4°C overnight, against A50 buffer containing 1 mM DTT and 0.5 mM ATP. After the addition of 0.5 mM fresh ATP to the dialyzed sample, it was centrifuged at 20,000 x g for 10 min at 4°C to remove aggregates and contaminating phosphorylated myosin. The purified protein was labeled with Alexa Fluor 546 NHS Ester (Invitrogen), snap-frozen in liquid nitrogen, and stored at –80°C.

### ZIP kinase

Recombinant human ZIPK was prepared according to a previous report with slight modifications (Kondo et al. 2012). Briefly, the protein was expressed in *E. coli* (Rosetta 2 (DE3), Merck Millipore) at 25°C for 1 h in the presence of 1 mM IPTG. The GST-tagged proteins were purified using a glutathione Sepharose column (GSTrap HP, GE Healthcare), followed by overnight dialysis against A50 buffer containing 1 mM DTT at 4°C. Samples were centrifuged at 627,000 x g for 20 min at 4°C to remove aggregates, snap-frozen in liquid nitrogen, and stored at –80°C.

### Arp2/3 complex

The Arp2/3 complex was purchased from Cytoskeleton Inc. The protein was diluted with a buffer containing 20 mM Tris-HCl (pH 7.5), 25 mM KCl, 1 mM MgCl_2_, 0.5 mM EDTA, 0.1 mM ATP, and 1 mM DTT. The protein was snap-frozen with liquid nitrogen and stored at –80°C.

### Profilin

Profilin was purified according to our previous report (Kubota et al. 2017).

### Capping protein

Capping protein was purified according to our previous report (Yamazaki et al. 2024).

### Cell culture and live imaging

MDCK cells (no. RCB0995: RIKEN Bioresource Center) stably expressing Lifeact-miRFP703 were a kind gift from Dr. Kazuhiro Aoki (Kyoto University). The cells were maintained in DMEM (08459-64, Nacalai Tesque) supplemented with 10% fetal bovine serum (FBS; F7524-500ML, Sigma), 100 units/mL penicillin and 100 μg/mL streptomycin (168-23191, Wako) in a 5% CO_2_ humidified incubator at 37°C. For the passage of the MDCK cells, the growth media was removed, and the cells were washed with 3 mL PBS. Next, 3 mL PBS containing 1 mM EDTA and 0.25% trypsin (18172-94: Nacalai Tesque) was added, immediately removed and the cells were incubated for 5 min at 37 °C. Prewarmed DMEM was added to resuspend the cells.

To observe the trypsinized cells, the cells were detached from the 6-well plates under dark conditions. Next, the cells were suspended with 200 μL FluoroBrite (A1896702, Gibco) supplemented with 10% FBS, 100 units/mL penicillin, and 100 μg/mL streptomycin, and plated on the collagen-coated 35 mm glass-bottom dishes. The imaging was started within 10 min after the seeding. During live-cell imaging, cells were incubated with a stage incubator set to 37 °C and containing 5% CO_2_ (STXG-IX3WX; Tokai Hit). The 488 nm laser was globally illuminated from the objective lens for the activation of iLID. The light intensity applied for the activation was approximately 0.1 μW/μm^2^.

### Transfection

Since the VCA domain of WAVE1 fused with fluorescence protein tends to aggregate for long-term expression, all of the experiments were performed by transient expression. The MDCK cells were electroporated by using a 4D-Nucleofector (Lonza) according to the manufacturer’s instructions (CA152 program) and attached solutions. For transient expression, 10 μg of total plasmids were mixed with the mass ratio of SspB:iLID = 1:4 to achieve efficient membrane translocation by light. After electroporation, the cells were incubated at room temperature for 10 min and plated on the 6-well culture plates (VTC-T6, AS ONE) or collagen-coated 35-mm glass-base dishes (D11130H, Matsunami). The cells were imaged 18-24 hours after the electroporation.

### Drug treatment

An Arp2/3 complex inhibitor, CK-666 (SML0006, Sigma-Aldrich), was diluted with DMSO (D2650, Sigma-Aldrich) at a concentration of 50 mM. The CK-666 treatment at a final concentration of 50 μM started 30 min before trypsinization. The concentration was maintained during live-cell imaging. A myosin II ATPase inhibitor, blebbistatin (B0560, Sigma-Aldrich), was diluted with DMSO (D2650, Sigma-Aldrich) at a concentration of 10 mM. The assembled SMM filaments were treated with blebbistatin at a final concentration of 50 μM 10 min before mixing with other proteins. The concentration was maintained during imaging. Although blebbistatin has been reported to be inactivated by blue light, the measured illumination intensity was at least 100-fold weaker than the intensity which cannot be inactivated (Fig. S4a, S4b) (Sakamoto et al. 2005).

### Confocal microscopy

An inverted microscope (ECLIPSE Ti2, Nikon) equipped with an EMCCD camera (iXon Life 888, ANDOR), a spinning disk confocal unit (CSU-W1, Yokogawa Electric Corporation), and an oil immersion objective lens (CFI Plan Lambda D 60x, N.A. 1.42, Nikon) was used. The excitation laser and fluorescence filter settings were as follows: Excitation laser, 405 nm, 488 nm, 561 nm, and 640 nm; dichroic mirror, DM 405/488/568/647; emission filters, 447/60 nm, 525/50 nm, 617/73 nm, and 685/40 nm. The microscope was controlled by NIS-elements software (Ver. 5.42.03, Nikon).

For global light illumination (Fig. 1, S1), 500 msec of pulsed blue light (488 nm) was illuminated through the objective lens with 20 sec intervals. For local light illumination (Fig. 2, 3, 4, 5, 6, S2, S3, S4, S5, S6, S7, S8), a digital micromirror device module (Nikon) and a light source SOLA (Lumencor) were mounted on the ECLIPSE Ti2 system and 500 msec of pulsed blue light was illuminated through the objective lens with 20 sec intervals. The filter settings for the digital micromirror device were as follows: Dichroic filter, 500; excitation filter 470/40 nm; emission filter, 535/50 (GFP-B, Nikon). To reduce the light intensity, an ND filter (NE220B, Thorlabs) was inserted into the light path of the SOLA. The light intensity irradiated by the digital micromirror device (DMD) was measured by a power meter (TQ8210, Advantest). Briefly, the sensor panel was attached to the objective lens, then blue light was illuminated and intensity was recorded with various light intensities and illumination patterns. The light intensity for inducing actin polymerization was maintained at 1 nW/μm^2^ without the ND filter, whereas with the ND filter, it was controlled from 0.04 to 0.18 nW/μm^2^. In all experiments using the DMD, the field of view of the microscope was 155 × 216 µm. Unless otherwise specified, within a single field of view: Fig. 2, 3, S2, S3, and S4 contained four ROIs of 37 × 37 µm each, Fig. 4, 6, and S6 contained ten ROIs of 25 × 25 µm each, and Fig. 5 contained six ROIs of 25 × 25 µm each, illuminated at varying intensities.

### TIRF microscopy

Single-molecule imaging was performed using hand-made TIRF microscopy. An inverted microscope (IX73, Olympus) equipped with an EMCCD camera (iXon3, Andor), the excitation laser (OBIS 488 nm LS, Coherent), and an oil immersion objective lens (PlanApo 60x, N.A. 1.45, TIRFM, Olympus) were used.

To obtain the diffusion coefficient of His-mEGFP-iLID on the SLB under the saturated density, His-ShadowG-iLID and His-mEGFP-iLID were mixed at a molecular ratio of 1000:1. After loading the protein mixture into the flowcell, we continuously illuminated excitation laser for 30 sec to reduce the weak background signal derived from His-ShadowG-iLID. Then, after the photobleaching of His-ShadowG-iLID, the bright signal derived from His-mEGFP-iLID was detected. Each particle trajectory over 1 sec was manually traced and then the mean-square displacement (MSD) was calculated. Based on the relationship MSD(Δ*t*) = 4*D*Δ*t* in lateral diffusion and using the slope of the linear approximation of the averaged MSD between molecules, the diffusion coefficient *D* of His-mEGFP-iLID was determined to be 0.38 μm^2^/sec.

### Confirmation of the binding of FMN to His-mEGFP-iLID

To confirm the requirement of FMN as a cofactor to the purified iLID, absorbance of 25 μM His-mEGFP-iLID at 450 nm with various FMN concentrations was measured using a UV-vis spectrometer (BioSpectrometer kinetic, Eppendorf). To subtract the background absorbance derived from His-mEGFP-iLID and FMN, 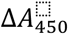 was calculated as 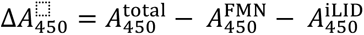, where 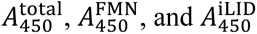 correspond to the 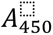 of 25 μM His-mEGFP-iLID containing FMN at each concentrations, 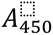 of FMN at each concentrations, and 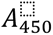 of 25 μM His-mEGFP-iLID, respectively.

### Preparation of the lipid bilayer

1-palmitoyl-2-oleoyl-sn-glycero-3-phosphocholine (POPC) and 1,2-dioleoyl-sn-glycero-3-[(N-(5-amino-1-carboxypentyl)iminodiacetic acid)succinyl] (nickel salt) (DGS-NTA(Ni)) were purchased from Avanti Polar Lipids. 1,2-Dihexadecanoyl-sn-Glycero-3-Phosphoethanolamine (Marina Blue DHPE) was purchased from Invitrogen. Each lipid was dissolved in chloroform within a round-bottom glass tube at a concentration of 20 mM. Next, DGS-NTA(Ni) was mixed with POPC at 2 mol% or 6 mol% in a new glass tube. To visualize the lipid, Marina Blue DHPE was also mixed at 2 mol% as needed. The lipids were dried by vacuuming overnight. The lipid-dried film was then hydrated with PBS (pH 6.5) overnight. The final lipid concentration was adjusted to 1 mM. The hydrated lipids were dispersed by five freeze-thaw cycles using liquid nitrogen and hot water at 30 °C. The lipids were stored at –30 °C and used within two months.

Functionalized coverslips were prepared as follows. First, coverslips (Matsunami, 24 × 60 mm, thickness No.1, non-coating) were sonicated in a mixture of 80% Ethanol and 5% NaOH for 90 min. Then, the coverslips were washed with Milli-Q water 5 times and sonicated again in Milli-Q water for 30 min. Finally, the coverslips were washed with Milli-Q water 5 times and stored in Milli- Q water at room temperature. The coverslips were used within 2 weeks. We then made a hand-made flow cell by putting double-sided tapes and then putting an 18 × 18 mm non-treated coverslip (Matsunami) on the functionalized coverslip. Before use, the functionalized coverslip was dried with a nitrogen gas stream and cleaned in the plasma cleaner (PDC-32G: Harrick Plasma) for 5 min. The volume of the flow cell was about 10 μL and the thickness was about 300 μm. The lipid solution was diluted 4 times with PBS (pH 6.5), filtrated using a Mini-Extruder Set (610000: Avanti Polar Lipids), and loaded into the flow cell. The flow cells were incubated for 10 min to make a lipid bilayer by vesicle fusion. Unfused vesicles were removed by washing with 50 μL of Milli-Q water.

### Preparation of the actin polymerization solution

All the following procedures were conducted at room temperature under red light to avoid activating iLID. First, 50 μL of His-mEGFP-iLID at a concentration of 7 μM was loaded into the flow cell. After 5 min incubation, unbound His-mEGFP-iLID was removed by washing with 50 μL of A50 buffer. Immediately after the washing, the actin polymerization solution was loaded into the flow cell, and the periphery of the flow cell was sealed with VaLaP. Then, the flow cell was mounted on the microscope and the imaging was started within 2 min.

During 5 min incubation of His-mEGFP-iLID, the actin polymerization solution was prepared. The total volume of the mixture was 20 μL in A50 buffer and the final composition was 2 mM Trolox, 10 μM FMN (F8399, Sigma-Aldrich), energy mix (1 mM ATP, 10 mM phosphocreatine, 0.1 mg/mL creatine phosphokinase, 10 mM DTT in A50 buffer), profilin, actin (10 mol % labeled with Alexa647), capping protein, Arp2/3 complex, and SspB-mScarlet-I-VCA. Note that actin was incubated on ice for more than 10 days after thawing and then used within 2 months. SMM and cofilin were also added to the solution, as needed. Although SMM and cofilin were detected by the same image channel as the SspB-mScarlet-I-VCA, it was easy to distinguish these proteins from SspB-mScarlet-I-VCA because of their distinct distributions. The concentrations of each protein are noted in each figure legend. To phosphorylate SMM, 1 μM SMM (40 mol % labeled with Alexa546), 100 nM ZIP kinase, and energy mix were mixed and incubated at 25 °C for 90 min. The SMM solution was used within 2 days.

### Pyrene actin polymerization assay

To measure the actin polymerization activity by pyrene fluorescence, 20 μM Actin (5 mol % labeled with pyrene) in a G-buffer was kept on ice more than 3 days before the measurement to completely disassemble oligomers contaminated in the freshly thawed G-actin solution from frozen stocks. First, 20 μL of 20 μM Actin was loaded into 6-wells of a 384-well plate. Next, 60 μL of 4/3× A50 buffer containing 4/3× energy mix and ABPs was loaded. The final concentrations were 2 mM Trolox, 10 μM FMN, 15 μM profilin, 25 nM capping protein, 50 nM Arp2/3 complex, 7 μM His-mEGFP-iLID, and 1 μM SspB-mScarlet-I-VCA. The fluorescence intensities of pyrene for each condition were simultaneously measured using a fluorescence plate reader (FluoroskanFL, Thermo Fisher Scientific) at 25°C for 20 min at 15 sec intervals. The minimum value for each condition was defined as the background noise and subtracted from each time-series data point.

### Measurement of SMM filament length

To induce SMM filament formation, 1 μM SMM (40 mol % labeled with Alexa546) was incubated with 100 nM ZIP kinase, energy mix in A50 buffer at 25 °C for 90 min. Then, the sample was diluted 100 times with A50 buffer containing 1 mM ATP and 10 mM DTT. The diluted sample was observed under confocal microscopy (see “confocal microscopy”). An oil immersion objective lens (CFI Plan Lambda D 100 x, N.A. 1.45; Nikon) was used. After background subtraction, the FWHM value along the short axis of SMM filaments was used to determine the threshold and then the raw images were binarised. The length of the long axis of each filament was measured as the filament length. The overlapping filaments were manually excluded from the analysis.

### Imaging analysis

Fluorescence imaging data were analyzed and quantified using Fiji (Image J). The fluorescence images were quantified after subtracting the background using the rolling ball function in Fiji unless otherwise described. The quantified data was plotted using Python. Three-dimensional images were reconstructed by the depth-coded alpha blending function of NIS-elements software (Nikon). To quantify the cortical fluorescence intensity of MDCK cells, we used a custom-made Fiji macro (Fig. 1c, 1d, 1g, S1b, S1c, S1d, S1f). Briefly, after the subtraction of the background, the images of Lifeact- miRFP703 were binarized. Next, to smooth the cell periphery, the images were enlarged twice and shrunk twice using dilate and erode functions in Fiji, respectively, and added as the first ROI. The first ROI was shrunk by 4 pixels and added as the second ROI. Finally, the area between the first and the second ROIs was analyzed as the cortical region, and the internal area of the second ROI was analyzed as the cytoplasmic region. The plot in Fig. 1e was generated using the same dataset as Fig. 1c, 1d. Briefly, for each cell, we quantified the translocation levels of SspB and Lifeact (F-actin) to the cortex. These values were normalized by the mean cortical intensity before light illumination. Additionally, we defined the mean cortical SspB intensity during t = 2–8 min as the SspB expression level. These three parameters were then plotted with a color code, allowing visualization of the relationship between SspB expression level, SspB translocation efficiency, and actin polymerization. To quantify the fluorescence intensity on the SLB, a single z-slice nearest to the coverslip surface was measured unless otherwise described. To measure the actin pillar length, the z-axis images were reconstructed by NIS-elements software (Fig. 3e, S5f). The distance from z = 0 µm to the center of the pillar was manually measured. To quantify the total actin intensity in the pillars (Fig. S4c, S5f), 3D slices were generated using Fiji. After the background subtraction, the integrated intensities in each ROI on the SLB were measured. To show the actin flow, the function of maximum intensity projection in Fiji was used (Fig. 5d, 5f, 5g, 5k). The temporal color-coded image of the actin flow was created after filling the floating protein aggregations with black color (Fig. 5h). To quantify the fluorescence intensity of Alexa647-actin in the presence of cofilin (Fig. 6b, S7a, S7b), the maximum intensity projection images were used because the actin pillars were detached from the SLB after stopping illumination. To quantify the actin disassembly level (Fig. 6c), the fluorescence intensities of actin 10 min after stopping illumination were divided by the fluorescence intensities of actin 10 min after starting illumination (the last frame before stopping illumination) in each pillar. The samples not showing visible actin polymerization were excluded from the scatter plots. Because the cofilin-induced actin disassembly levels changed exponentially with initial actin density in both our experiments and previous report (Manhart et al. 2019), the nonlinear least squares fitting approach using a power-law model of SciPy was applied. For the fitting, the initial actin fluorescence greater than 0.3 to eliminate the influence of the background signal.

### Statistics and reproducibility

In all box-and-whisker plots, the box shows the quartiles of data with the whiskers denoting the minimum and maximum except for the outliers detected by 1.5 times the interquartile range. The statistical analysis was performed by Python. The experiments were repeated at least twice independently to confirm reproducibility. The sample number for quantification is listed in the figure legends.

## Acknowledgments

We thank all members of the Miyazaki Laboratory for their helpful discussions and assistance. We are deeply grateful to Dr. Linh T. Tran for her generous support with the additional experiments. This work was supported by JSPS KAKENHI (Grant number 22J00094, 23K14179 to K.Y.; 21K19220, 22H05171 to M.M.), JST PRESTO (Grant number JPMJPR20ED to M.M.), RIKEN Incentive Research Project to K.Y. and RIKEN special postdoctoral researcher funding to K.Y. We appreciate Dr. François Robin and Dr. Kazuhiro Aoki for their fruitful advice and discussion.

## Author contributions

K.Y. and M.M. designed the research. K.Y. performed experiments. K.Y. analyzed data. K.Y. and M.M. wrote the manuscript.

## Supplementary Note

### Estimation of the actin network density from fluorescence images

To estimate the density of the actin networks assembled by illumination, we first measured the fluorescence intensity of homogeneous F-actin solutions with known concentrations. Briefly, 90 μM actin (10 mol % labeled with Alexa647) was polymerized in a polymerization buffer containing 50 mM KCl, 1 mM MgCl_2_, 1 mM EGTA, 20 mM 2-mercaptoethanol, 1 mg/mL BSA, 1 mM ATP, and 20 mM HEPES (pH7.2) at 25 °C for 30 min <u>(Shrivastava et al. 2015)</u>. During polymerization, the SLB was formed in a hand-made flow cell. To minimize the effect of the background signal from out of focus, we made a thin flow cell using two pieces of plastic wrap as a spacer. The thickness measured by a microscope was 12.75 ± 2.25 μm, which was comparable to the height of actin structures assembled by illumination. The polymerized actin solution was serially diluted with the polymerization buffer and loaded into the flow cells. The fluorescence intensities of F-actin solutions 4 μm above the glass surface were measured and plotted (Fig. 6d, S8a).

Next, we compared the fluorescence intensities of F-actin solutions with light-induced actin networks. We simply compared the raw fluorescence intensities of the F-actin solutions and pillars without subtracting the background noise. This is because the rolling ball subtraction method is not suitable for nearly homogeneous images, such as those of F-actin solutions. The concentrations of the actin networks in the sparse conditions, the dense conditions with 0.05-0.15% outputs, and the dense conditions with 0.15-0.45% outputs are estimated to be 0-20, 20-40, and 40-60 μM, respectively.

Assuming that the concentration of the densest network is ∼50 μM, the density of actin molecules is 30,000/μm^3^. It can be reasonably approximated that actin filaments are uniformly distributed in the pillar and extend vertically from the SLB <u>(Yamazaki et al. 2024)</u>. Given that an actin filament elongates by 2.7 nm in length with each binding of actin monomers <u>(Kubota et al.</u> <u>2018)</u>, the number of actin molecules in each filament is ∼370 molecules per μm along the Z-axis. Therefore, the density of actin filaments in the XY plane is estimated to be ∼81 filaments/μm^2^. This corresponds to 9 filaments per μm, meaning that the filaments are spaced ∼110 nm apart on average. Applying the same estimation method to the actin networks at a density of 25 μM, which allows myosin penetration into the network (Fig. 5a, S8a), results in ∼160 nm apart on average. Similarly, at a density of 15 μM in the sparse condition, the spacing is estimated to be ∼250 nm on average.

### Estimation of the actin filament length

The length of branched actin filaments is determined by the competitive binding of actin monomers and capping protein (CP) to the barbed end of an elongating filament. Therefore, the length distribution of branched filaments can be estimated from the association rate constants of ATP-actin and capping protein to the barbed end. The probability of the binding of CP to a filament after the polymerization of 𝑛 actin molecules can be expressed as

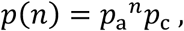

where the binding probabilities of actin monomer and CP are given as 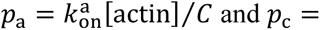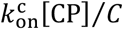, respectively, and 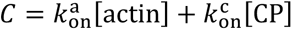 is the normalization constant. By substituting the literature values (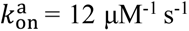 <u>(Pollard 2016)</u>, 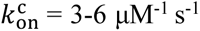 <u>(Schafer et al. 1996; Wear et al. 2003)</u>) and the concentrations used for the experiments ([actin] = 5 μM, [CP] = 0.025 μM), the expected value of 𝑛 is calculated as

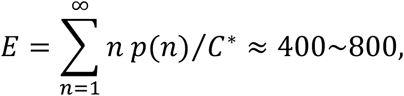

where 𝐶^∗^ is the normalization constant represented as

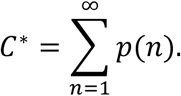

Given that an actin filament elongates by 2.7 nm with each binding of actin monomers <u>(Kubota et al.</u> <u>2018)</u>, the expected filament length is approximately 1-2 μm. Note that the filament length was estimated to be ∼300 nm in the actin pillars assembled from similar protein compositions to our experiments <u>(Bieling et al. 2016)</u>. Given that the concentration of capping protein is 4 times lower in our experiments, the potential filament length estimated from the previous study would be ∼1,200 nm. This value is also consistent with our estimation.

## Supplementary Figures

**Figure S1.**
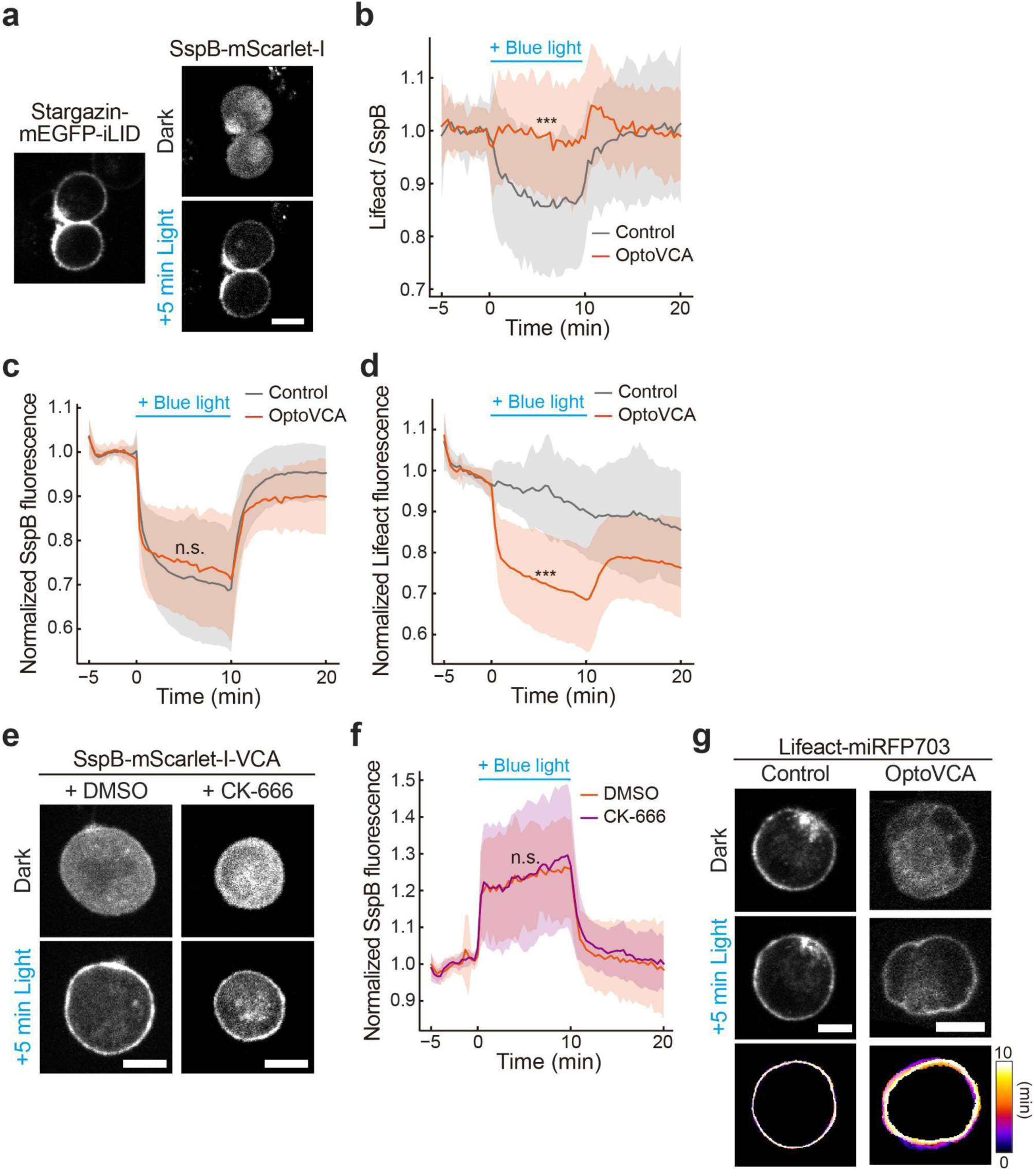
OptoVCA system for *in vivo* experiments. (a) Representative images of MDCK cells expressing Stargazin-mEGFP-iLID (left) and SspB-mScarlet-I (right). Upon blue light illumination, SspB-mScarlet-I translocated from the cytoplasm to the plasma membrane. (b) The ratio of normalized cortical Lifeact to normalized cortical SspB in Fig. 1c, d. The mean values (bold lines) are plotted as a function of time with the SD. *n* = 37 cells. (c) The normalized cytoplasmic fluorescence intensity of SspB-mScarlet-I (Control) or SspB-mScarlet-I-VCA (OptoVCA). The mean values (bold lines) are plotted as a function of time with the SD. *n* = 37 cells. (d) The normalized cytoplasmic fluorescence intensity of Lifeact-miRFP703. The mean values (bold lines) are plotted as a function of time with the SD. *n* = 37 cells. (e) Representative images of the cells expressing SspB-mScarlet-I-VCA treated with DMSO (left) or CK-666 (right) before (upper) and after (lower) blue light illumination. (f) Normalized cortical fluorescence intensity of SspB-mScarlet-I-VCA treated with DMSO or CK-666. The mean values (bold lines) are plotted as a function of time with the SD. *n* = 18 and 17 for DMSO and CK-666 treated cells, respectively. (g) Comparison of shape changes between control cell (left) and OptoVCA expressing cells (right). Temporal color-coded images show the cell edges after starting illumination. All scale bars, 20 μm. *p* values were calculated by unpaired two-sided *t*-test at *t* = 5 min. ***, *p* < 0.001. n.s., *p* ≥ 0.05.

**Figure S2.**
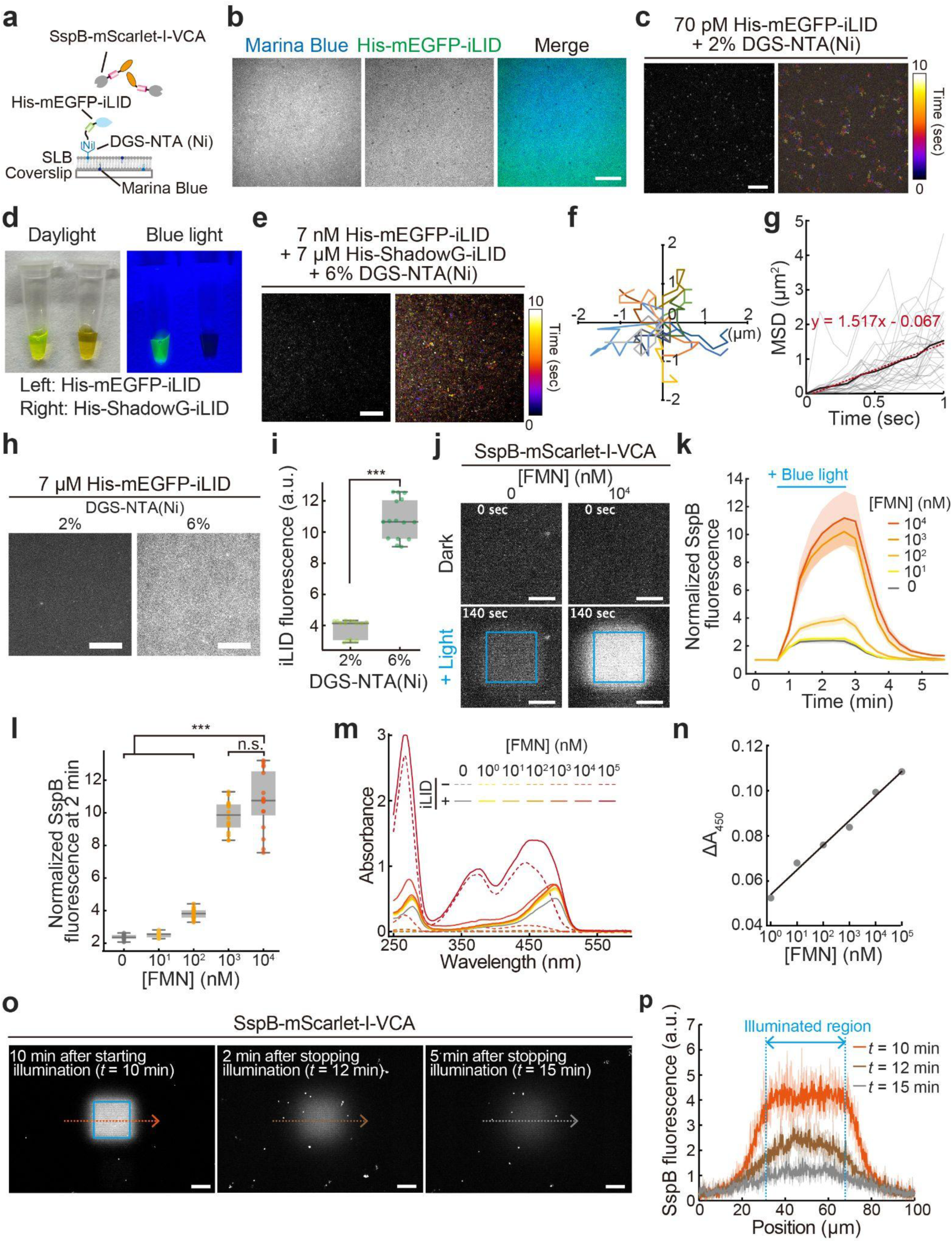
Optimization of iLID-SspB system for *in vitro* experiments. (a) Schematic illustration of the OptoVCA system without cytoskeletal proteins. (b) Representative images of SLB visualized by 2% Marina Blue lipid. 6% DGS-NTA(Ni) was added to POPC to anchor His-mEGFP-iLID on SLB. (c) TIRF images of lipid-bound His-mEGFP-iLID. Left, a single snapshot. Right, a temporal color-coded image reconstructed from 100 images taken over 10 sec. (d) Purified His-mEGFP-iLID and His-ShadowG-iLID under the daylight (left) and blue light (right). (e) TIRF images of lipid-bound His-mEGFP-iLID mixed with His-ShadowG-iLID. Left, a single snapshot. Right, a temporal color-coded image reconstructed from 100 images taken over 10 sec. (f) Representative trajectories of 15 His-mEGFP-iLID molecules over 1 sec. (g) Mean-square displacement (MSD) analysis of His-mEGFP-iLID. Black line, red dashed line, and gray lines indicate averaged MSD, linear approximation, and 30 single MSDs, respectively. (h) Representative images of His-mEGFP-iLID on SLB containing 2% (left) or 6% (right) DGS-NTA(Ni). (i) Fluorescence intensities of His-mEGFP-iLID in (h). *n* = 15 for each sample. *p* values were calculated by unpaired two-sided *t*-test. ***, *p* < 0.001. (j) Representative images showing the accumulation of SspB-mScarlet-I-VCA on SLB before (upper) and after (lower), and without FMN (left) and with 10^4^ nM of FMN. Condition: 150 nM SspB-mScarlet-I-VCA. Blue-squared regions were illuminated with 1 nW/μm^2^. (k) Fluorescence intensities of SspB-mScarlet-I-VCA in the presence of various concentrations of FMN. The mean values (bold lines) are plotted as a function of time with the SD. (l) Fluorescence intensities of SspB-mScarlet-I-VCA at *t* = 2 min in (k). *n* = 16 for each sample. Statistical significance was determined using one-way ANOVA followed by Tukey’s multiple comparisons test. ***, *p* < 0.001. n.s., *p* ≥ 0.05. (m) Absorption spectra of FMN in the presence or absence of 25 μM His-mEGFP-iLID. (n) Confirmation of the binding of FMN to His-mEGFP-iLID detected by the absorbance at 450 nm. (o) Whole microscopic field of view showing recruitment and dissociation of SspB-mScarlet-I-VCA after starting and stopping illumination at each indicated time point. Blue-squared region was illuminated with 1 nW/μm^2^. Condition: 150 nM SspB-mScarlet-I-VCA. (p) Line profiles of SspB-mScarlet-I-VCA intensity along the dashed arrows in (o). The bold lines indicate the mean values of three independent experiments (thin lines). All scale bars, 20 μm.

**Figure S3.**
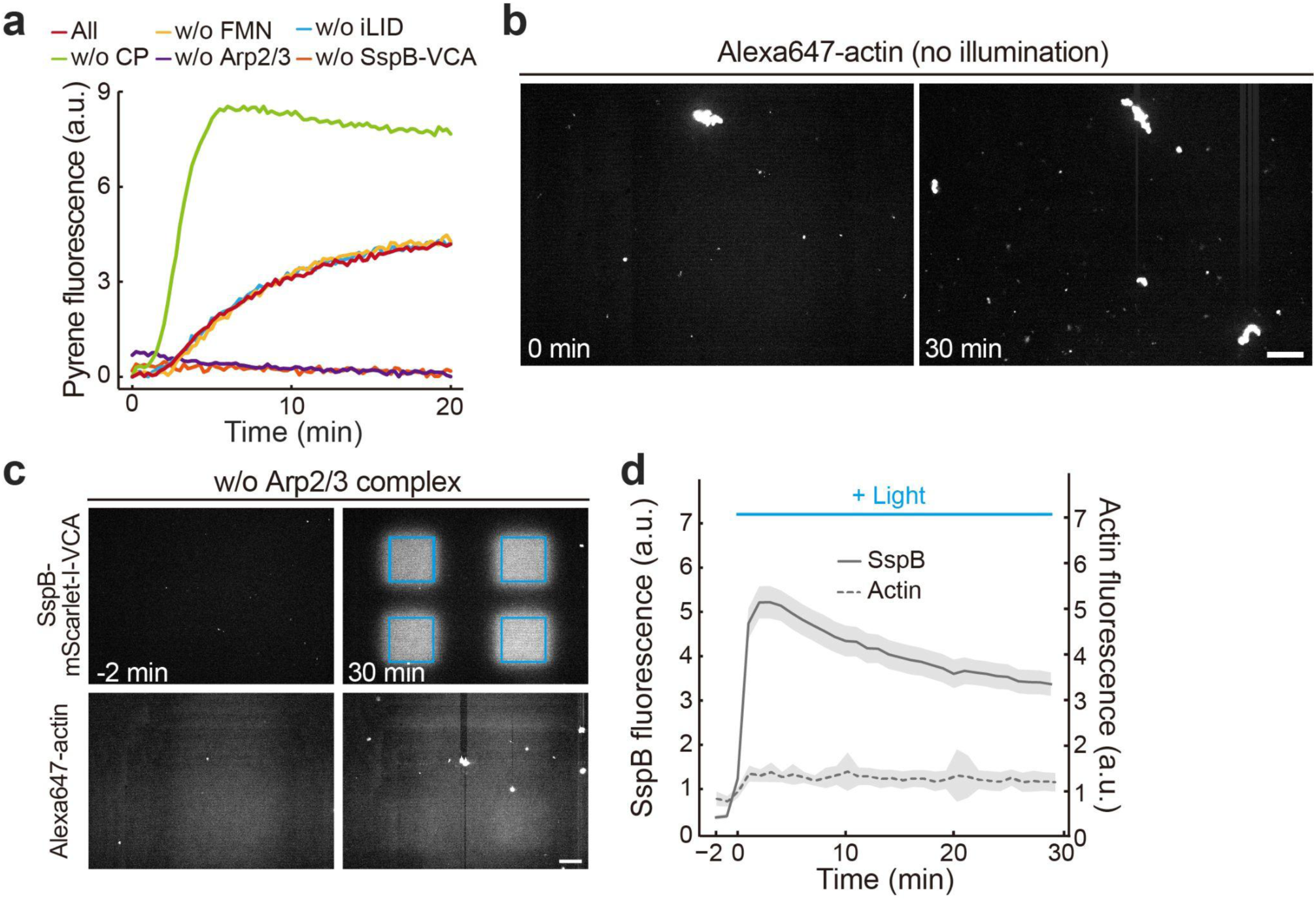
Optimization and validation of *in vitro* OptoVCA system. (a) Pyrene fluorescence assay using all components or omitting each component to confirm actin polymerization activity. (b) Representative images of Alexa647-actin under the dark condition. Condition: 5 μM Actin, 15 μM Profilin, 100 nM Arp2/3 complex, 25 nM CP, and 150 nM SspB-mScarlet-I-VCA. (c) Whole microscopic field of view showing SspB-mScarlet-I-VCA (upper panels) and Alexa647-actin (lower panels) in the absence of Arp2/3 complex at each indicated time point. Blue-squared regions were illuminated with 1 nW/μm^2^. (d) Fluorescence intensities of SspB-mScarlet-I-VCA (solid line) and Alexa647-actin (dashed line) in (c). The mean values (bold lines) are plotted as a function of time with the SD. *n* = 12 for each condition. All scale bars, 20 μm.

**Figure S4.**
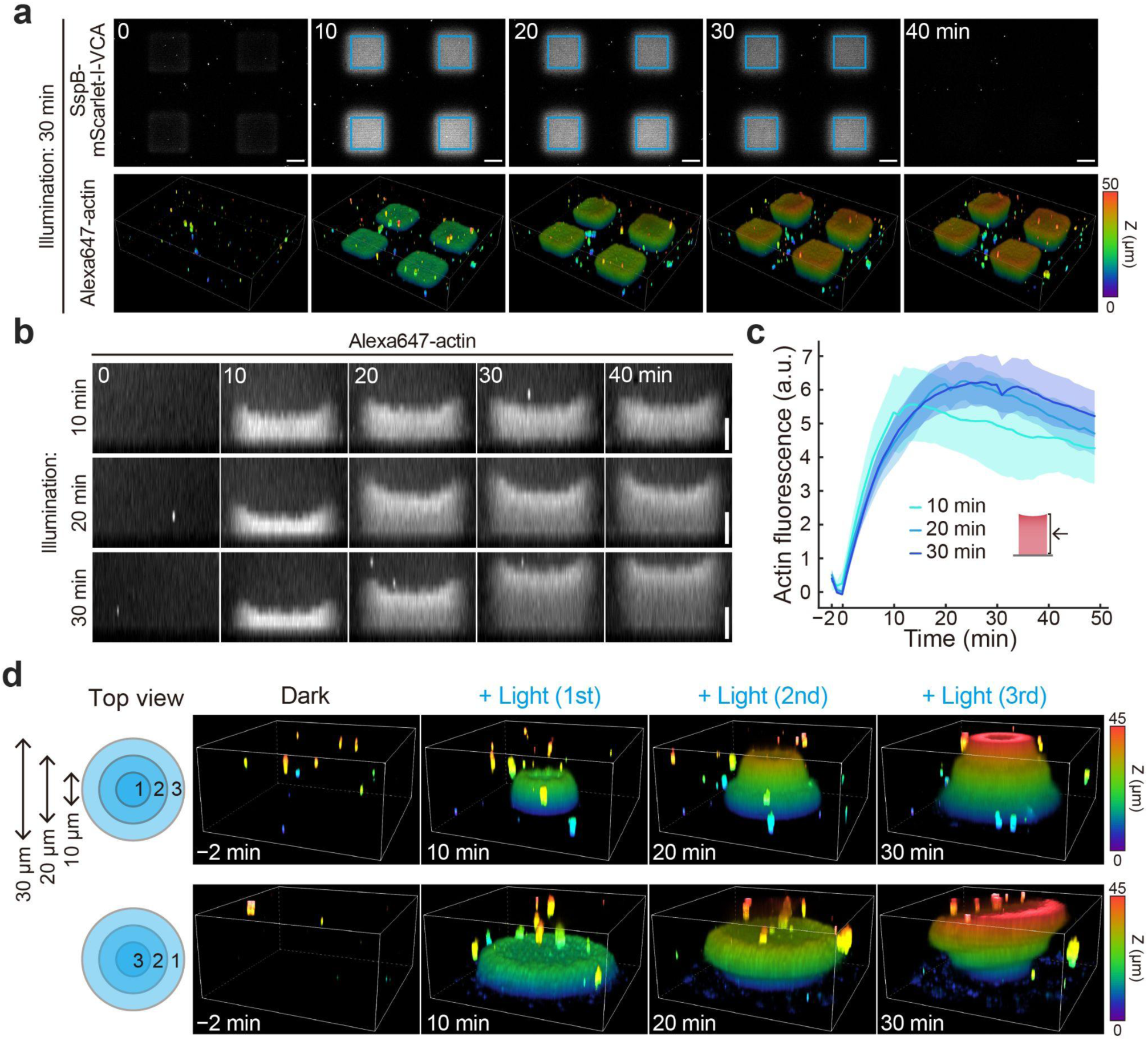
Temporal control of actin network assembly. (a) Whole microscopic field of view showing SspB-mScarlet-I-VCA (upper panels) and Alexa647-actin (lower panels) in the absence of Arp2/3 complex at each indicated time point. Blue-squared regions were illuminated with 1 nW/μm^2^ for 30 min. (b) Three-dimensional slices showing the growth dynamics of actin pillars at each indicated illumination time. (c) Total actin intensity in the slice image of (b). The mean values (bold lines) are plotted as a function of time with the SD. *n* = 12 for each condition. (d) Schematic illustration of the three-step illumination process (left) and representative images of the created actin structures (right). Each indicated region was illuminated for 10 min in the order specified by the numbers in the illustration. Condition: 5 μM Actin, 15 μM Profilin, 100 nM Arp2/3 complex, 25 nM CP, and 150 nM SspB-mScarlet-I-VCA. All scale bars, 20 μm.

**Figure S5.**
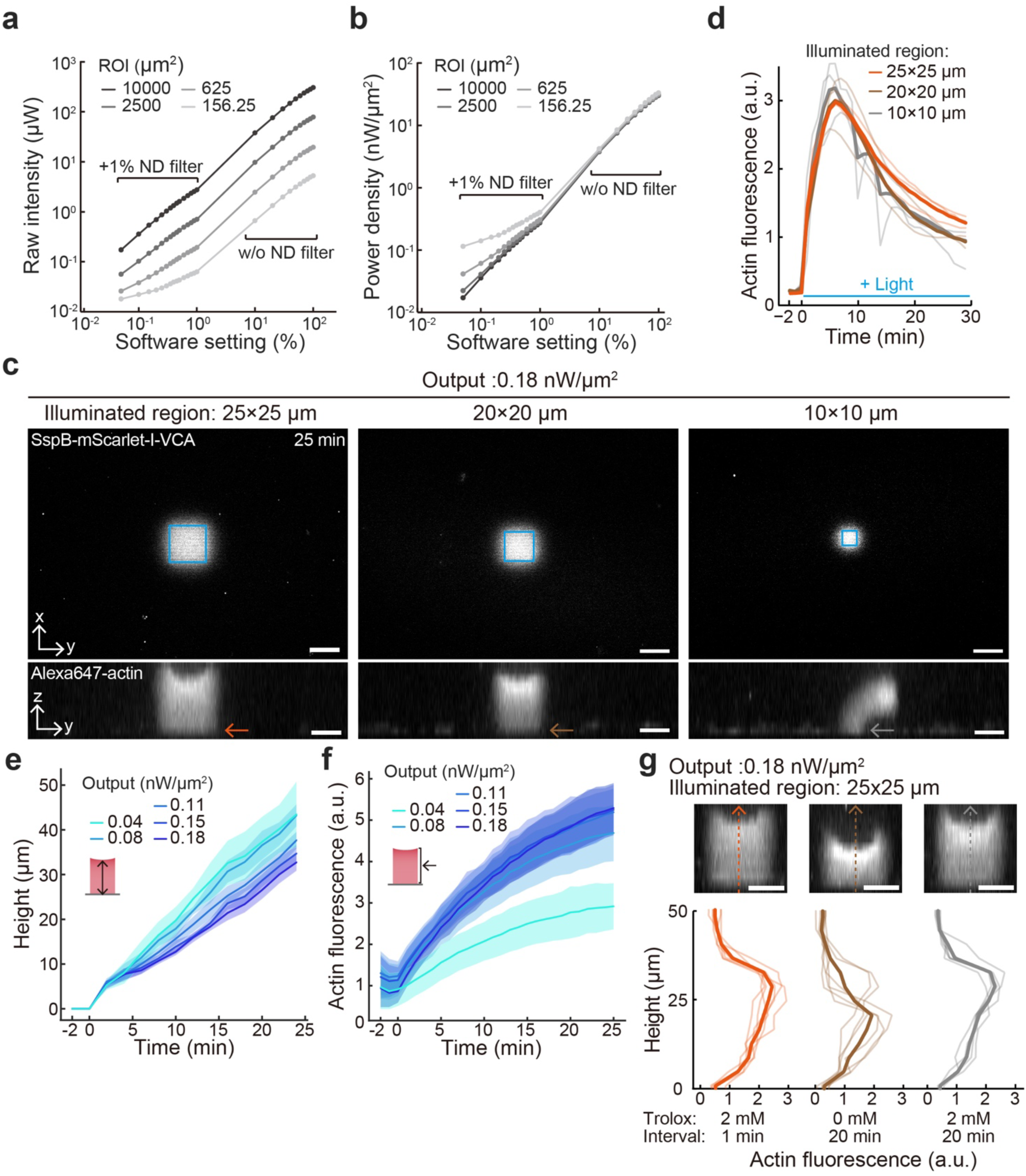
Control of actin density by light intensity. (a, b) Raw light intensities with different-sized illumination patterns (a) and the power densities normalized by the illuminated areas (b) plotted as a function of the output power of the DMD light source. (c) Whole microscopic field of view showing recruitment of SspB-mScarlet-I-VCA to the single center region with three different ROI sizes (upper panels) and the formed actin pillar 25 min after starting illumination (lower panels). Blue-squared regions were illuminated. (d) Fluorescence intensities of Alexa647-actin in (c). The bottom plane of the actin pillar was used for the quantification (Indicated as arrows in (c)). The bold line indicates the mean values of three independent experiments (thin lines). (e, f) Growth dynamics (e) and total fluorescence intensity (f) of the actin pillars in each indicated illumination output in Fig. 4. The mean values (bold lines) are plotted as a function of time with the SD. *n* = 6 for each condition. (g) Line profiles of Alexa647-actin fluorescence intensity in each indicated condition along the dashed arrows in upper panels. Blue light was illuminated for 20 min. The bold line indicates the mean values of 6 independent experiments (thin lines). (c, g) Condition: 5 μM Actin, 15 μM Profilin, 100 nM Arp2/3 complex, 25 nM CP, and 150 nM SspB-mScarlet-I-VCA. All scale bars, 20 μm.

**Figure S6.**
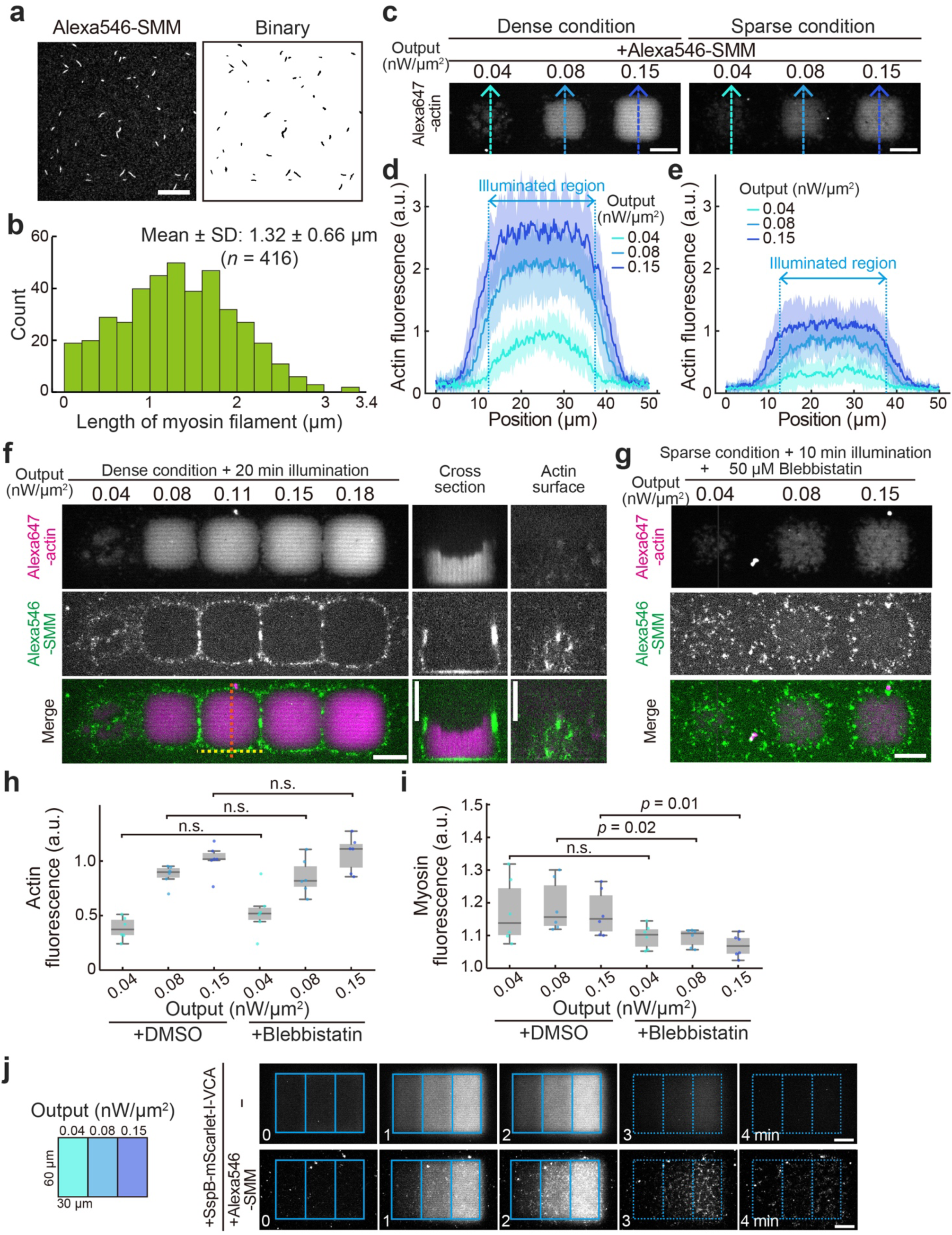
Effects of actin network density on myosin. (a) Myosin filaments labeled with Alexa546 (left) and its binarized image (right). Scale bar, 10 μm. (b) Histogram showing the length distribution of myosin filaments. (c) Representative images of polymerized actin in the presence of 200 nM Alexa546-SMM. Dense condition: 5 μM Actin, 15 μM Profilin, 100 nM Arp2/3 complex, 25 nM CP, and 150 nM SspB-mScarlet-I-VCA. Sparse condition: 2 μM Actin, 6 μM Profilin, 100 nM Arp2/3 complex, 10 nM CP, and 150 nM SspB-mScarlet-I-VCA. Images taken after 5 min illumination are shown. All scale bars, 20 μm. (d, e) Line profiles of polymerized actin in dense condition (d) and sparse condition (e). A single plane located 4 μm above the lipid membrane was quantified. Dashed arrows in (c) indicate scanned lines. The mean values (bold lines) are plotted as a function of time with the SD. *n* = 10 for each condition. (f) Spatial distribution of SMM in dense conditions. The cross-section and actin surface images correspond to the orange and yellow dashed lines in the merged image, respectively. Condition: 5 μM Actin, 15 μM Profilin, 100 nM Arp2/3 complex, 25 nM CP, 150 nM SspB-mScarlet-I-VCA, and 200 nM Alexa546-SMM. All scale bars, 20 μm. (g) Spatial distribution of SMM treated with 50 μM blebbistatin in the sparse condition. Condition: 2 μM Actin, 6 μM Profilin, 100 nM Arp2/3 complex, 10 nM CP, 150 nM SspB-mScarlet-I-VCA, and 200 nM Alexa546-SMM. Scale bar, 20 μm. (h, i) Fluorescence intensities of actin (h) and myosin (i) in the sparse condition. *p* values were calculated by unpaired two-sided *t*-test. n.s., *p* ≥ 0.05. *n* = 6 for each sample. (j) Illumination pattern to induce actin flow (left illustration) and temporal dynamics of the recruited SspB-mScarlet-I-VCA in the absence (upper panels) or presence (lower panels) of 200 μM Alexa546-SMM. Blue light was illuminated for 2 min from *t* = 0 to 2 min. The illuminated regions were indicated as blue boxes and dashed blue boxes during and after the illumination, respectively. Scale bars, 20 μm.

**Figure S7.**
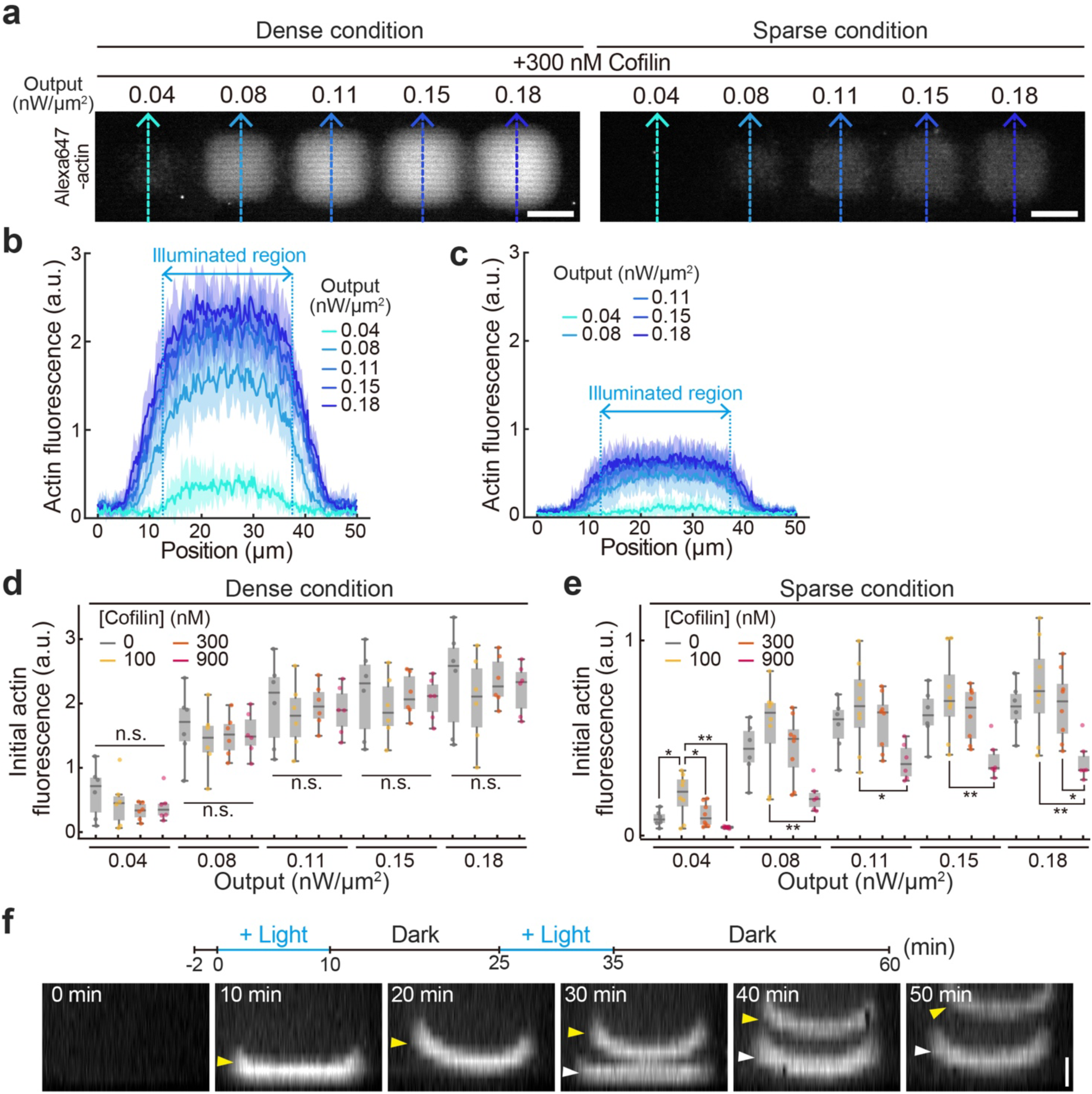
Effects of actin network density on cofilin. (a) Representative images of polymerized actin in the presence of 300 nM Cofilin. Dense condition: 5 μM Actin, 15 μM Profilin, 100 nM Arp2/3 complex, 25 nM CP, and 150 nM SspB-mScarlet-I-VCA. Sparse condition: 2 μM Actin, 6 μM Profilin, 50 nM Arp2/3 complex, 10 nM CP, and 150 nM SspB-mScarlet-I-VCA. Images taken after 10 min illumination are shown. (b, c) Line profiles of polymerized actin in dense condition (d) and sparse condition (e). A single plane located 4 μm above the lipid membrane was quantified. Dashed arrows in (a) indicate scanned lines. The mean values (bold lines) are plotted as a function of time with the SD. *n* = 6 and 8 for dense and sparse conditions, respectively. (d, e) Initial actin fluorescence intensities in the dense (d) and sparse (e) conditions after 10 min illumination. In the dense conditions, *n* = 6 for each cofilin concentration. In the sparse conditions, *n* = 6, 8, 8, 6 for 0, 100, 300, 900 nM cofilin, respectively. Statistical significance was determined using one-way ANOVA followed by Tukey’s multiple comparisons test. **, *p* < 0.01. *, *p* < 0.05. n.s., *p* ≥ 0.05. (f) Three-dimensional raw images showing the floating actin networks corresponding to Fig. 6h. Yellow and white arrowheads indicate the first and second polymerization, respectively. All scale bars, 20 μm.

**Figure S8.**
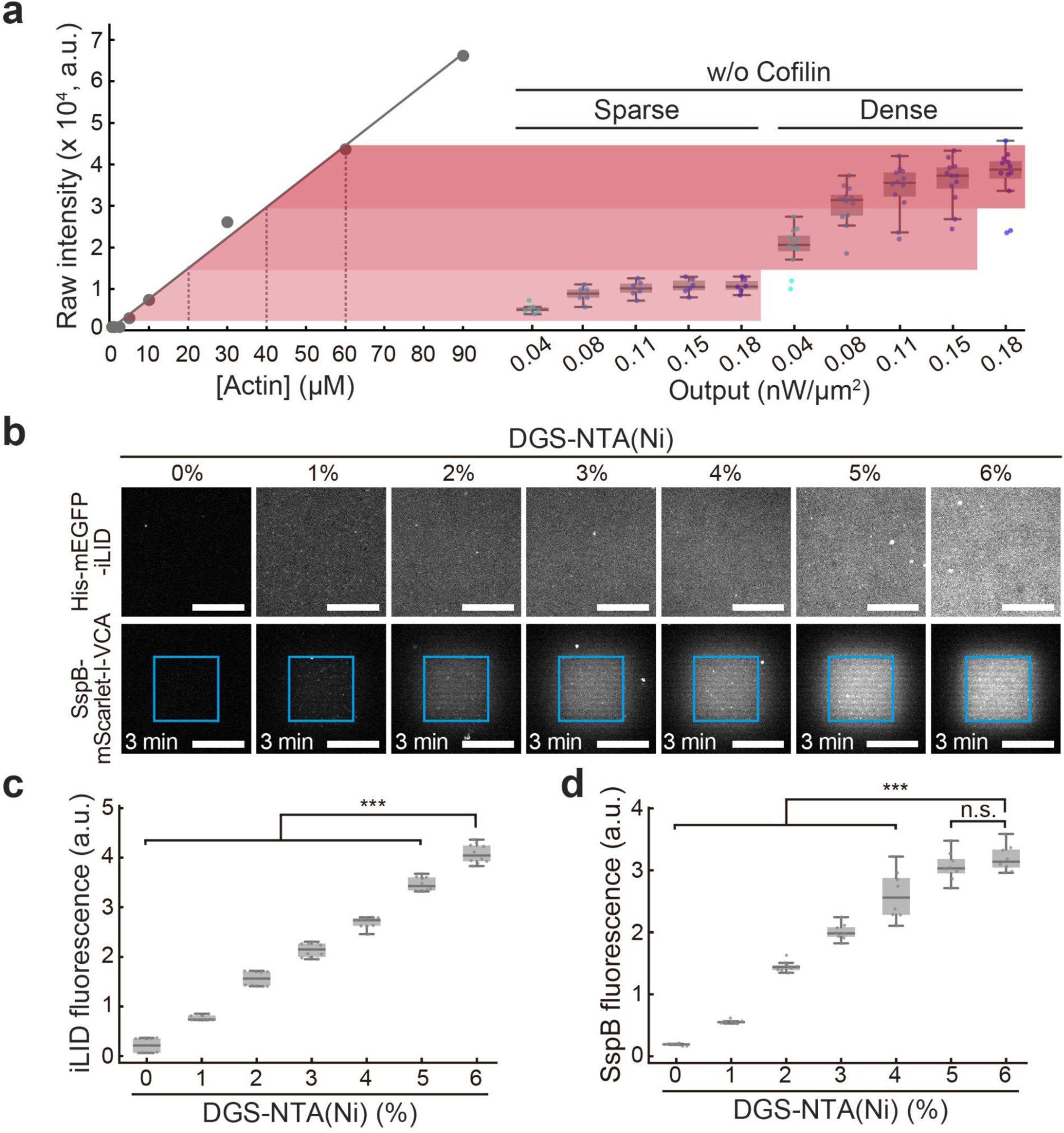
Estimation of actin network density and VCA density. (a) Comparison of the raw fluorescence intensities between homogeneous F-actin solutions (left) and light-induced actin networks (right). *n* = 6 and 12 for sparse and dense conditions, respectively. (b) Representative images showing His-mEGFP-iLID (upper panels) and recruitment of SspB-mScarlet-I-VCA (lower panels) on the SLB with various DGS-NTA(Ni) lipid densities. Blue-squared regions were illuminated with 1 nW/μm^2^. All scale bars, 20 μm. (c) Fluorescence intensities of His-mEGFP-iLID in (b). *n* = 10 for each sample. (d) Fluorescence intensities of SspB-mScarlet-I-VCA in the illuminated region in (b). Intensities 3 min after starting illumination were quantified. *n* = 8 for each sample. Statistical significance was determined using one-way ANOVA followed by Tukey’s multiple comparisons test. ***, *p* < 0.001. n.s., *p* ≥ 0.05.

## Supplementary Movie legends

**Movie S1.** Growth dynamics of actin network seeds upon blue light illumination. The blue rectangular region indicates the illuminated area. Scale bar, 20 μm.

**Movie S2.** Growth dynamics of actin networks induced by three-step illumination patterns.

**Movie S3.** Growth dynamics of actin pillars assembled by 10 min (left), 20 min (middle), and 30 min (right) illumination. Scale bars, 20 μm.

**Movie S4.** Growth dynamics of actin pillars induced by five different illumination intensities.

**Movie S5.** Formation of an actin sheet with a density gradient, followed by SMM-driven actin flow toward the dense region. Blue rectangular regions indicate illuminated areas. Scale bar, 20 μm.

**Movie S6.** Formation of an actin sheet with a density gradient, followed by SMM-driven actin flow from both sides toward the central dense region. Blue rectangular regions indicate illuminated areas. Scale bar, 20 μm.

**Movie S7.** Floating actin networks induced by various illumination patterns.

**Movie S8.** Floating actin networks induced by repetitive illuminations.

